# Exoproteome of calorie-restricted humans identifies complement deactivation as an immunometabolic checkpoint reducing inflammaging

**DOI:** 10.1101/2025.08.04.668533

**Authors:** Manish Mishra, Hee-Hoon Kim, Yun-Hee Youm, Elsie Gonzalez-Hurtado, Konstantin Zaitsev, Tamara Dlugos, Irina Shchukina, Christy Gliniak, Eric Ravussin, Subhasis Mohanty, Albert C. Shaw, Philipp E. Scherer, Maxim N. Artyomov, Vishwa Deep Dixit

## Abstract

Caloric restriction (CR) extends lifespan, yet the convergent immunometabolic mechanism of healthspan remains unclear. Using longitudinal plasma proteomics analyses in humans achieving 14% CR for 2 years, we identified that inhibition of the complement pathway is linked to lower inflammaging. The protein C3a (and its cleaved form) was significantly lowered by CR, thus reducing inflammation emanating from three canonical complement pathways. Interestingly, circulating C3a levels are increased during aging in mice, with visceral adipose tissue macrophages as the predominant source. In macrophages, C3a signaling via ERK elevated inflammatory cytokine production, suggesting the existence of an autocrine loop that promotes inflammaging. Notably, long-lived FGF21-overexpressing mice and PLA2G7-deficient mice exhibited lower C3a in aging. Specific small molecule-mediated systemic C3 inhibition reduced inflammaging, improved metabolic homeostasis, and enhanced healthspan of aged mice. Collectively, our findings reveal that complement C3 deactivation is a metabolically regulated inflammaging checkpoint that can be harnessed to extend healthspan.

## INTRODUCTION

Aging is featured by the accumulation of damage and the chronic activation of the innate immune system that leads to time-dependent functional decline at the molecular, cellular, tissue, and organ levels, resulting in a compromised healthspan^1,2^. Therefore, identification of causal mechanisms that perturb homeostasis in aging and their successful targeting through dietary or pharmacological means are needed to promote the health of older adults and stave off chronic diseases. Calorie restriction (CR), a non-pharmacological intervention, has emerged as the most effective approach to extending the healthspan and lifespan of an organism^3,4^. Notably, the Comprehensive Assessment of Long-term Effects of Reducing Intake of Energy (CALERIE-II) clinical trial showed that CR boosts the thymic and metabolic function while lowering the inflammation in humans^5,6^. However, the precise pathways and related mechanisms underlying the beneficial effects of CR warrant further investigation.

The protective innate immune response of the host is central to the organism’s survival. Throughout evolution, the complement system is documented as one of the highly conserved and ancient groups of proteins across different species, ranging from invertebrates to humans, making it a key component of the protective innate response^7^. In addition to the documented role of the complement in opsonization and cell lysis, growing evidence implicates that complement proteins as important regulators of cellular functions, such as controlling tissue homeostasis^8–10^. Complement system activation, a cascade of more than 50 soluble or membrane-bound glycoproteins in humans, mainly involves three pathways: classical, lectin, and alternative^11^. Importantly, complement activation, irrespective of the pathways involved, converges at the central component, complement 3 (C3). C3 is cleaved and activated by C3 convertase, producing byproducts C3a and C3b, which regulate inflammation^12^. Chronic inflammation, one of the major hallmarks of aging^13^, drives most of the age-associated complications, suggesting the potential contribution of C3 and C3a. An elevated C3 level is negatively correlated with longevity^14^.

Longevity is extended with CR^15,16^. Many studies have recently investigated the role of complement proteins in mammalian aging and reported elevated C3 concentrations in serum and organs of aged mice^17^. Notably, the elevation of C3 and its activated products, such as C3a, is a common signature observed in aging^18,19^. In support of a strong association between aging and innate immune activation, recent proteomic analysis of human organ samples spanning approximately 50 years of age also identified complement and NLRP3 inflammasome activation as the top upregulated pathways^20^. Although recent reports have investigated the importance of the classical pathway protein C1q in aging-related phenotypes^21,22^, the overall C3 levels derived from the activated lectin and the alternative pathway have been largely overlooked. Considering the biological source and the relevance of elevated complement activation, such as C3a, seen during aging, remains incompletely understood. We investigated C3 and its downstream inflammatory effector C3a, which account for the entire complement pathways implicated. In addition to the liver as a primary synthesis site^23^, C3 is locally produced by adipocytes and macrophages^24,25^. Macrophage activation is well documented as a driver of age-associated inflammation or inflammaging^26^. However, the contributing role and involved mechanisms of macrophages in age-associated C3 regulation are still open questions.

In this study, utilizing unbiased plasma proteomics, we discovered that 2 years of 14% CR reduced multiple complement proteins, including C3a, and related pathways in middle-aged healthy individuals, independent of body mass index (BMI). In addition, we found that CR slowed the adipose tissue proteomic aging clock. In contrast, in male and female mice, we observed that C3a expression increased with age, and this activation is predominantly derived from visceral adipose tissue (VAT). Specifically, adipose tissue macrophages (ATMs) were identified as the primary cellular source of age-associated C3 production, and downstream signaling through the extracellular signal-regulated kinase (ERK) pathway mediated the production of inflammatory cytokines in an autocrine manner. Accordingly, pharmacological inhibition of C3 improved healthspan and restrained inflammaging in aged mice. Moreover, we observed that the age-associated increase in C3a was partially blunted in long-lived fibroblast growth factor 21 (FGF21) transgenic mice and CR-mimetic phospholipase A2 group VII (PLA2G7) knockout mice compared to wild-type controls. Together, our reverse translational approach suggests that the inhibition of the complement system is a promising, clinically relevant target to reduce inflammaging and prolong healthspan.

## RESULTS

### CR rewires the directionality of the human exoproteome towards longevity pathways

To investigate the impact of two years of mild CR on the exoproteome of middle-aged healthy individuals, we performed an unbiased proteomics analysis using the Somascan 7K assay on longitudinal plasma samples from 42 CALERIE participants from the Pennington Biomedical Research Center cohort in Baton Rouge, collected at baseline and after 2 years of 14% CR (Fig. 1a). Most participants in this cohort were in their thirties and forties and were not obese (BMI <30), and they lost around 10% of their body weight after 2 years of CR (Fig. 1b and Extended Data Table 1). We detected a total of 7,029 proteins (SOMAmers) that passed quality control, and 262 proteins failed quality control (Fig. 1c and Extended Data Table 2). Principal component analysis showed that CR led to considerable changes in the plasma proteome of the participants (Extended Data Fig. 1a).

**Fig. 1.**
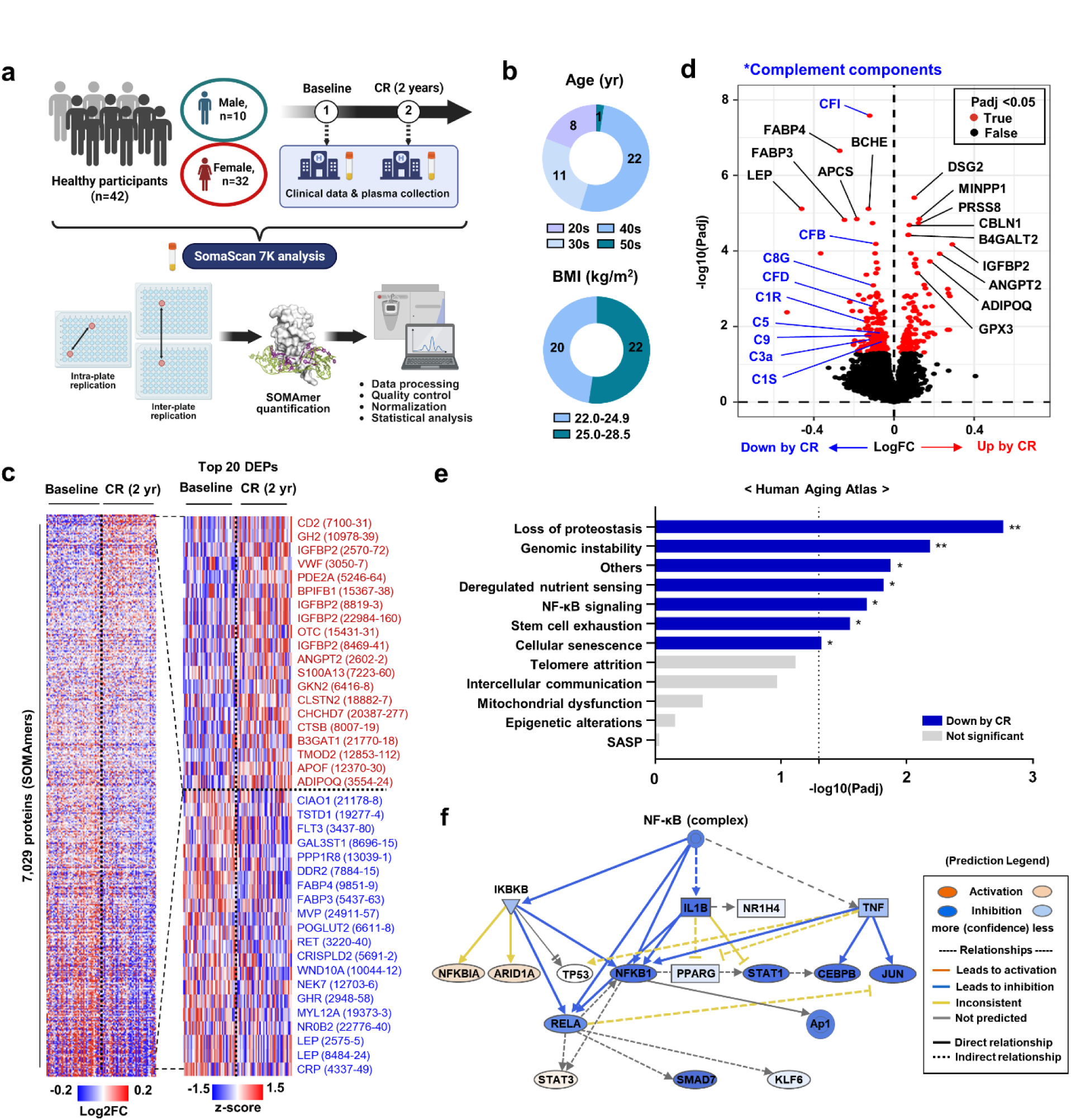
CR rewires the directionality of the human exoproteome towards longevity pathways. (a-f) SomaScan 7K proteomics was performed in the plasma of healthy individuals at baseline and 2 years after 14% caloric restriction (CR) (*n* = 42). (a) Schematic diagram of the overall study design. (b) Age (top) and BMI (bottom) of participants. (c) Heatmaps for all detected proteins (*n* = 7,029; left) and the top 20 increased or decreased differentially expressed proteins (DEPs) (right). (d) A volcano plot for the plasma proteomics analysis. (e) CR-associated changes in 12 hallmarks of aging pathways were analyzed, and a scoring analysis of each pathway was performed. *padj <0.05 and **padj <0.01. (f) The prediction of upstream regulator analysis for nuclear factor-kappa B (NF-κB) in Ingenuity Pathway Analysis. Most of the NF-κB signaling-related molecules were reduced by CR.

To explore the CR-induced proteomic shifts in detail, we performed differential expression analysis. We found that insulin-like growth factor-binding protein 2 (IGFBP2) was one of the top increased proteins after CR, while other IGFBPs remained unchanged except for IGFBP5, which was reduced (Fig. 1c,d and Extended Data Fig. 1b-h). IGFBP2 and IGFBP5 may impact mechanisms of aging by lowering the IGF-1 bioavailability and are related to reduced mortality^27^ and cellular senescence^28^, respectively. Although the circulating IGF-1 concentration upon CR in our cohort was not different, these findings confirm the importance of reduced IGF-1 bioavailability, a mechanism identified from mice^29^, for some beneficial effects of CR on aging in humans. As expected, CR significantly induced adiponectin levels but reduced leptin, fatty acid binding protein (FABP) 3 and 4^30^, and growth hormone receptor levels, all of which are indicative of healthy aging (Fig. 1c,d and Extended Data Fig. 1i-m)^31–34^. Of note, even though the participants were healthy and did not have inflammatory conditions, CR significantly decreased protein levels of complement components as well as other pro-inflammatory markers, such as c-reactive protein, tumor necrosis factor (TNF)-α, and C-C motif ligand 1 (CCL1) but did not affect interleukin (IL)-1β and IL-6 levels in the plasma (Fig. 1d and Extended Data Fig. 1n-r).

A recent study has established the Aging Atlas database, encompassing most of the hallmark pathways of aging based on previously published omics datasets^35^. We queried this database to examine the effects of CR on each aging hallmark pathway at the protein level. Interestingly, we found that none of the 12 aging hallmark pathways were increased, but more than half, including loss of proteostasis, genomic instability, deregulated nutrient sensing, nuclear factor (NF)-κB signaling, stem cell exhaustion, cellular senescence, and other aging-associated features, were significantly reduced by CR (Fig. 1e and Extended Data Fig. 1s). In particular, the prediction of upstream regulator analysis by Ingenuity Pathway Analysis (IPA) revealed that most of the NF-κB-regulated molecules were suppressed by CR (Fig. 1f), suggesting that plasma proteomics can reflect intracellular signaling hubs. These longitudinal analyses over 2 years of CR intervention in healthy individuals suggest reversal of some key aging hallmark signatures, particularly targeting both immune and metabolic pathways.

### CR in humans slows adipose tissue proteomic aging clock

We next investigated the target organ(s) that may contribute to exoproteomic rewiring of aging hallmark pathways in humans post CR. To examine this, we utilized a recently developed algorithm, ‘organage’, which enables the estimation of organ-specific age using human plasma proteomics data^36^. This model first maps plasma proteins to putative organ sources based on organ-enriched expression features, defined as being more than four times higher in the organ of interest than in any other organ, in the Genotype-Tissue Expression (GTEx) project. It then uses these organ-enriched proteins to model organ-specific age and measures the gap between the chronological age and predicted age (Fig. 2a). Interestingly, based on these analyses, we found that CR only significantly reduced the age gap in adipose tissue but had no effects on the lung, kidney, artery, muscle, and heart, and increased the age gap in the pancreas and intestine (Fig. 2b-i and Extended Data Fig. 2a-h).

**Fig. 2.**
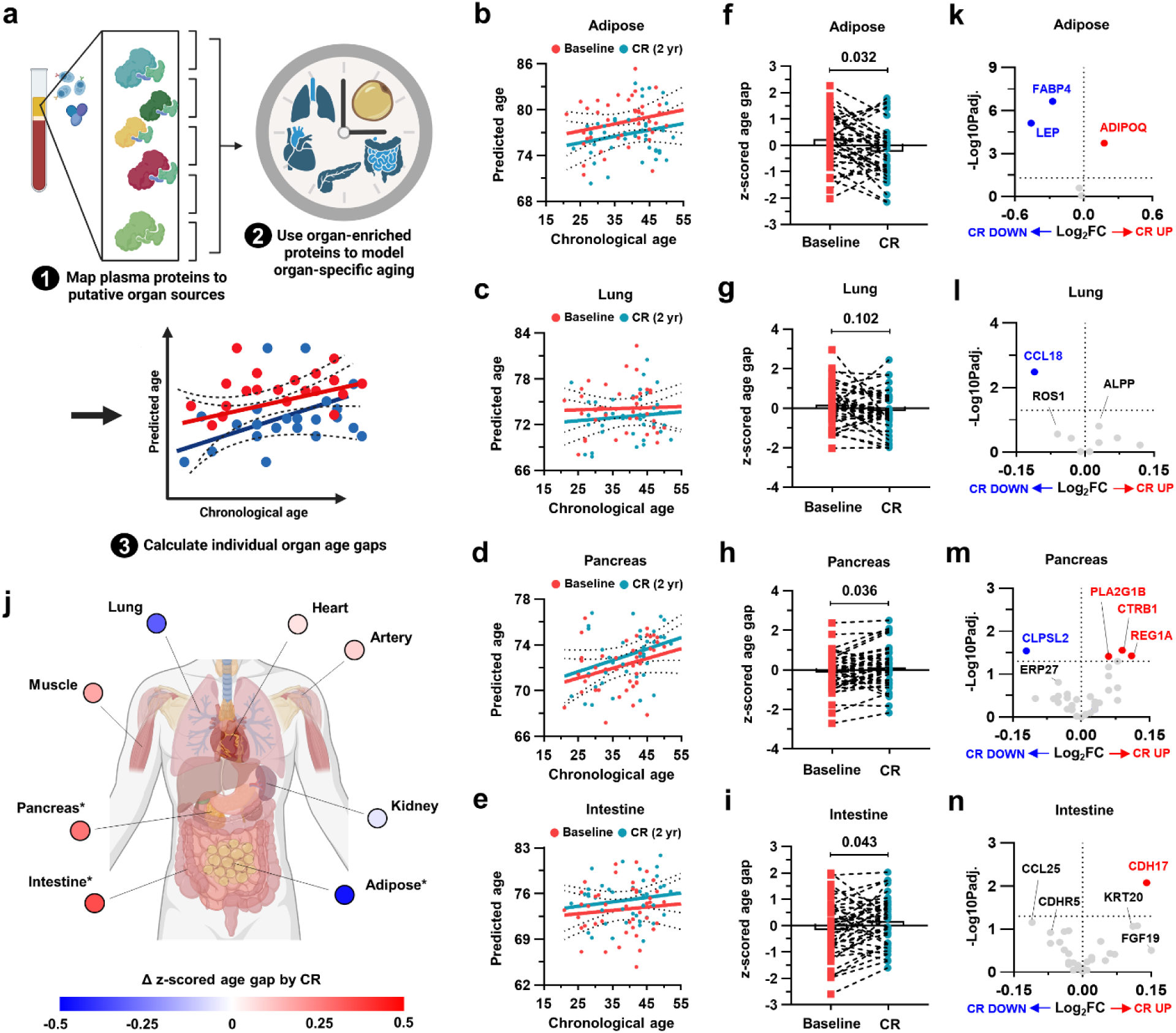
CR in humans slows adipose tissue proteomic aging clock. (a-n) Organ-specific aging signatures were analyzed based on the plasma proteome. (a) Schematic diagram for the analysis pipeline. (b-e) Linear regression analysis was performed to compare the predicted proteomic age before and after CR in adipose tissue (b), lung (c), pancreas (d), and intestine (e). (f-i) Organ age gap was analyzed in adipose tissue (f), lung (g), pancreas (h), and intestine (i). (j) Summary of the changes in the delta z-scored organ age gap by CR. (k-n) Volcano plots for the organ-specific proteins in adipose tissue (k), lung (l), pancreas (m), and intestine (n). Paired two-tailed t-tests were performed, and exact adjusted p-values were presented.

Since this algorithm predicts organ age based on the association of protein expression with age, rather than a causal relationship, we plotted all the organ-enriched proteins to examine the actual effects of CR on organ aging. For adipose tissue, we confirmed that CR beneficially modulated the adipose proteome, as shown in the differential expression analysis, with increased adiponectin and decreased leptin and FABP4 levels (Fig. 2k). Similarly, in the lung, CR significantly reduced CCL18 (Fig. 2l), which has been implicated in pulmonary fibrosis^37^. Notably, although the estimation of proteomic age showed that the age gaps in the pancreas and intestine increased after CR, the significantly increased pancreas-enriched or intestine-enriched proteins after CR have been known to play tissue-protective roles, such as lipid digestion (PLA2G1B), protection from pancreatitis (CTRB1)^38^ and inflammatory bowel disease (REG1A)^39^, and supporting intestinal barrier function (CDH17)^40^ (Fig. 2m,n). In addition, the kidney, artery, and muscle did not show any significantly regulated proteins, and BMP10, important in cardiac development and function^41^, was increased in the heart after CR (Extended Data Fig. 2i-l). The absence of change based on these in silico analyses may not fully capture the biology due to the lower sensitivities of detected protein features. Collectively, these data may suggest that CR is beneficial across various organs, with the most notable effects observed in the adipose tissue.

### CR downregulates C3a level independently of BMI

To explore the specific mechanisms driving the benefits of CR, we performed pathway enrichment analysis with the plasma proteomics data using the Gene Ontology (GO) and Kyoto Encyclopedia of Genes and Genomes (KEGG) database^42,43^. We found that CR upregulated pathways related to neuronal development, cardiac function, and tissue remodeling (Fig. 3a). In addition, CR suppressed pathways involved in the regulation of lipid storage, positive regulation of IL-6 production, and positive regulation of immune response (Fig. 3a). These results confirmed the previously reported immunometabolic advantages of CR^44–47^. Notably, the top downregulated pathways after CR were the ‘complement and coagulation cascades’ (Fig. 3a,b). Given that the expression levels of complement components increase with age and are implicated in various age-associated dysfunction^21,48,49^, we delved deeper into CR-induced changes of each complement component.

**Fig. 3.**
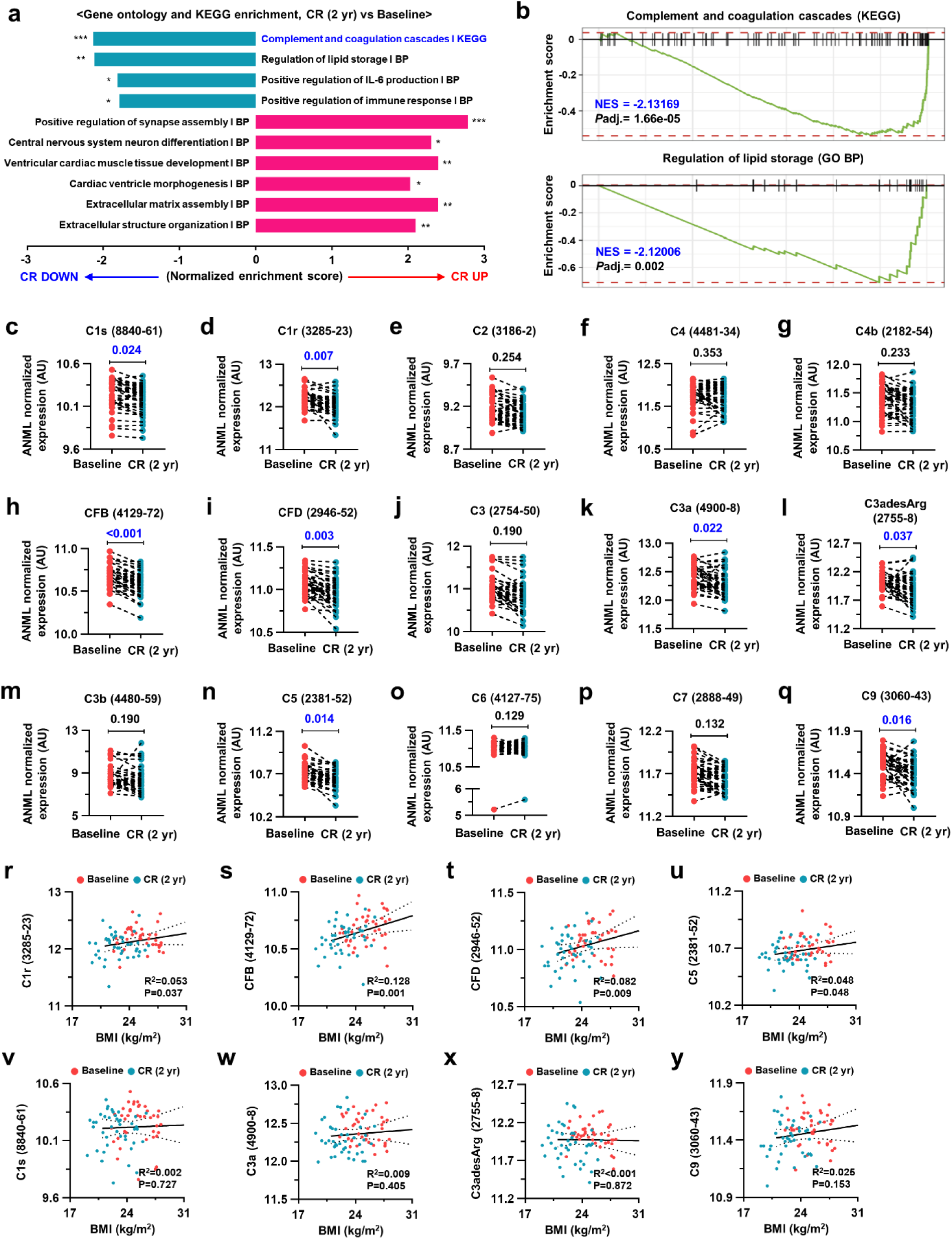
CR downregulates C3a level independently of BMI. (a) Gene set enrichment analysis for databases gene ontology (GO) (BP – biological process) and KEGG. Only significant pathways (adjusted p-value <0.05) were shown. (b) Enrichment plots for complement and coagulation cascades from the KEGG database and regulation of lipid storage from GO BP. (c-q) ANML normalized expression values for the indicated complement system components. (r-y) Linear regression analysis was conducted to examine the correlation between BMI and indicated complement components. For Figure 3c-q, paired two-tailed t-tests were performed, and adjusted p-values were presented.

We found that CR significantly reduced plasma levels of some early classical pathway-related components, such as C1s and C1r, but not C2, C4, and C4b (Fig. 3c-g). In addition, CR also suppressed the expression levels of alternative pathway-related components, specifically CFB and CFD (Fig. 3h,i). A central event during complement activation is the cleavage of C3 into C3a and C3b, which further promotes inflammation and the formation of the membrane attack complex (MAC), respectively. Although C3 and C3b levels were not affected, both C3a and C3adesArg, a more stable form of C3a with the c-terminal arginine cleaved, were significantly reduced by CR (Fig. 3j-m). Furthermore, among the MAC components, C5 and C9 showed significant reductions after CR, while C6 and C7 levels remained unchanged, and C8 was not detected (Fig. 3n-q). These results suggest that CR may suppress overall complement cascades rather than a specific pathway (Extended Data Fig. 3).

Since BMI has been reported to be associated with widespread changes in the circulating proteome in humans^50^, we examined the association between BMI and the 8 complement components (C1s, C1r, CFB, CFD, C3a, C3adesArg, C5, and C9) that were significantly reduced after CR. Of note, although the degrees of association were not strong, C1r, CFB, CFD, and C5 showed significant positive correlations with BMI, indicating that the reduced expression levels of these molecules could be derived from weight loss after CR (Fig. 3r-u). In contrast, C1s, C3a, C3adesArg, and C9 exhibited BMI-independent expression patterns (Fig. 3v-y). Impressively, these data indicate that the inhibition of C3a might be a unique CR mimetic that can be harnessed to extend immune and metabolic healthspan.

### Age-associated increase in C3a expression is unique to the VAT in mice

Given the data from the proteomics analysis of the CALERIE clinical trial, we utilized a reverse translational approach to further examine the source and roles of C3a during aging. First, we measured serum C3 levels in humans and mice to investigate whether aging per se could affect its production. Cross-sectional analysis of young (age 21-30) and older (age ≥65) adults demonstrated a significant increase in C3 concentrations in older adults, which appeared to be driven by elevated levels, particularly in females (Fig. 4a-c). Furthermore, analyses of C57BL/6N mice from the National Institute on Aging (NIA) and C57BL/6J from Yale rodent colonies revealed that aging was associated with a significant elevation of C3a in both sexes of mice, while C3 did not change (Fig. 4e and Extended Data Fig. 4a,b).

**Fig. 4.**
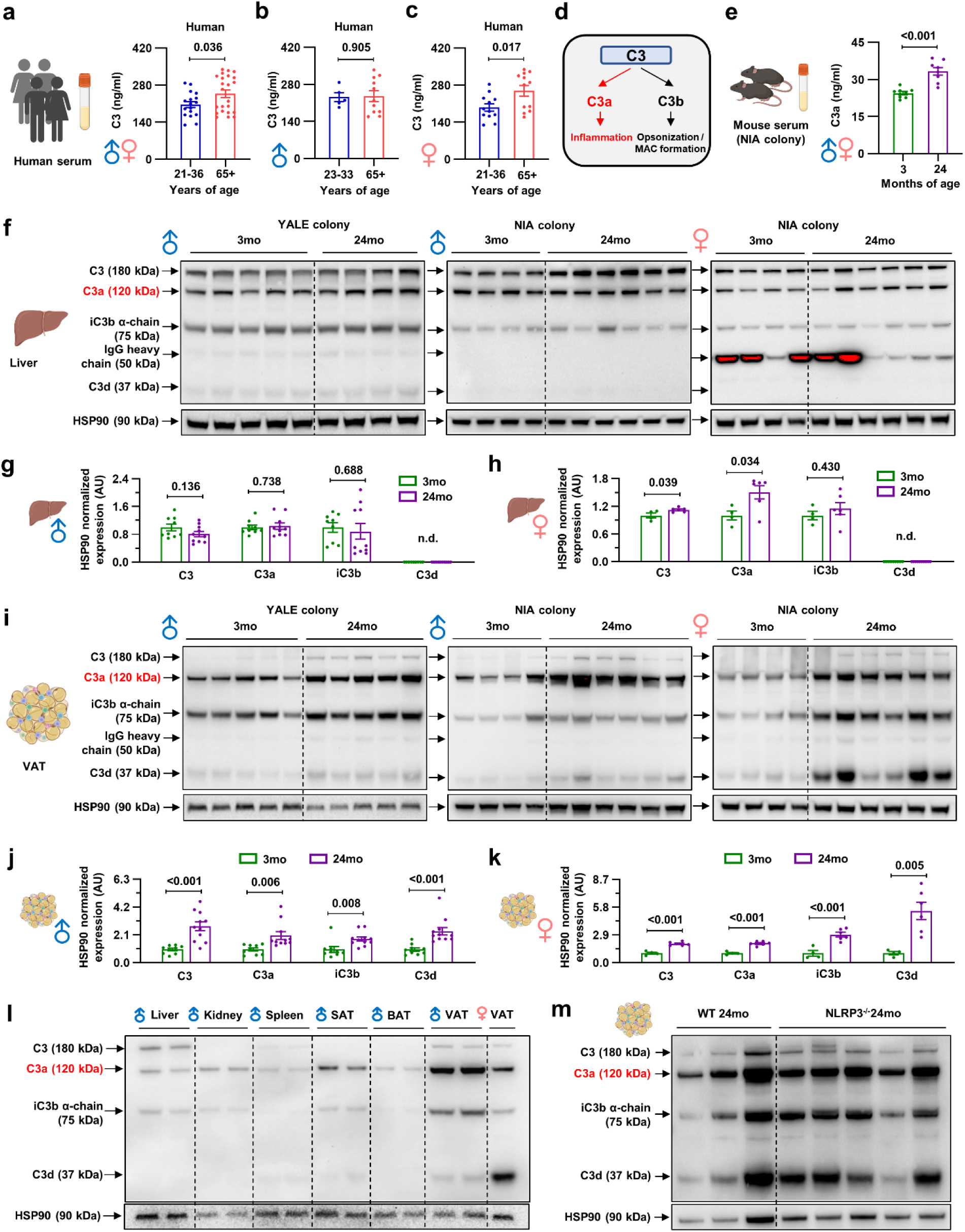
Age-associated increase in C3a expression is unique to the VAT in mice. (a-c) Human serum samples were analyzed for C3 ELISA in both sexes (a; *n* = 17 for 21-36 years old and *n* = 23 for >65 years old), males (b; *n* = 5 for 23-33 years old and *n* = 11 for >65 years old), or females (c; *n* = 12 for 21-36 years old and *n* = 12 for >65 years old). (d) Schematic diagram for C3 cleavage. (e) Mouse serum samples were analyzed for C3a ELISA in both sexes (*n* = 9 for 3-month-old and *n* = 8 for 24-month-old). (f-h) C3 western blot analysis of mouse liver. (f) Representative blots for C3. (g) Densitometry analysis of male mice (*n* = 9 for 3-month-old and *n* = 10 for 24-month-old). (h) Densitometry analysis of female mice (*n* = 4 for 3-month-old and *n* = 6 for 24-month-old). (i-k) C3 western blot analysis of mouse visceral adipose tissue (VAT). (i) Representative blots for C3. (j) Densitometry analysis of male mice (*n* = 9 for 3-month-old and *n* = 11 for 24-month-old). (k) Densitometry analysis of female mice (*n* = 4 for 3-month-old and *n* = 6 for 24-month-old). (l) Representative blots for C3 in the indicated tissues of 24-month-old mice. (m) Representative blots for C3 in 24-month-old WT (*n* = 3) and NLRP3^-/-^ (*n* = 5) mice. Unpaired two-tailed t-tests were performed, and exact p-values were presented. Data presented as mean±SEM.

Moving further, we investigated the principal source of increased serum C3a levels observed in aged mouse serum through biochemical assays of various organs, including the liver, visceral adipose tissue (VAT), subcutaneous adipose tissue (SAT), brown adipose tissue (BAT), spleen, and kidney from aged mice at Yale (C57BL/6J) and from NIA. We were able to detect either C3 (180 kDa) or its cleaved forms, such as C3a/C3α (120 kDa), iC3b α-chain (75 kDa), or C3d (37 kDa), in all tissues and samples analyzed. It has been previously reported that the liver is a primary organ contributing to serum C3 levels^51^. However, we observed only a modest increase in C3 and C3a liver expression levels in females, while male mice did not show any changes in C3 and its cleaved molecules with age (Fig. 4f-h). This suggests that the liver may not be a major source of the aging-associated elevation of serum C3a levels. Similarly, although there were alterations in the levels of C3 and/or its cleaved molecules in some tissues, we could not find an increase in C3a levels in the SAT, BAT, spleen, and kidney in either sex during aging (Extended Data Fig. 4c-j). Of note, we found that the age-associated increase in C3a levels was unique to the VAT in both sexes (Fig. 4i-k). In addition, the relative expression levels of C3a were the highest in the VAT among the analyzed tissues (Fig. 4l). Moreover, by analyzing in-house C57BL/6J male mice (maintained in a different location from the NIA colony), we consistently observed that the VAT, and not the liver, was the major source of C3a during aging (Fig. 4f,i), suggesting that these results were not dependent on animal facility or husbandry-associated microbiota. Given that VAT is a major source of age-associated C3a elevation and previous reports linking the NOD-, LRR-, and pyrin domain-containing protein 3 (NLRP3) inflammasome with complement proteins^52,53^, we asked whether the aged VAT-specific C3a elevation requires the NLRP3 inflammasome activation. We observed comparable expression of either C3 or its cleaved products in VAT from 24-month-old NLRP3-deficient mice and wild-type controls (Fig. 4m and Extended Data Fig. 4k), concluding that the unique expression of C3a in aged-VAT was NLRP3-independent. Together with the CALERIE proteomics data, which showed decreased proteomic age only in adipose tissue, these results suggest that VAT plays a pivotal role in the age-associated immune-metabolic dysregulations, possibly by producing C3a.

### Autocrine C3a signaling in adipose tissue macrophages regulates inflammaging

We next asked which complement pathway is involved in the elevation of C3a levels in the VAT of aged mice? By western blot analysis, we did not detect an increase in C1QA (classical pathway) or complement factor D (alternative pathway) during aging, and C2 (lectin pathway) was not detected in the VAT (Fig. 5a). Nevertheless, the ERK pathway, one of the downstream signaling pathways of the C3a receptor (C3AR1), was significantly activated with age in the VAT (Fig. 5a). Since previous studies have reported C3 production by both adipocytes^24^ and macrophages^25,54^, we next sought to identify the cellular source of C3a within the VAT. To do this, we isolated the mature adipocytes and the stromal vascular fraction (SVF), which contain various cell types, including immune cells, from the VAT of 3- and 24-month-old mice. In the mature adipocyte fraction, we could only detect C3d, but not C3 or C3a, and C3d was increased with age (Fig. 5b and Extended Data Fig. 5a). Notably, we observed all forms of C3 and its cleaved products in the SVF, where C3a and C3d were significantly increased with age (Fig. 5c and Extended Data Fig. 5b). Moreover, although we did not find age-associated changes in C1QA and CFD in whole VAT, there was a tendency for an increase in both proteins in the SVF with age, along with ERK activation (Fig. 5c and Extended Data Fig. 5b). These results suggest a non-adipocyte origin of C3a during aging.

**Fig. 5.**
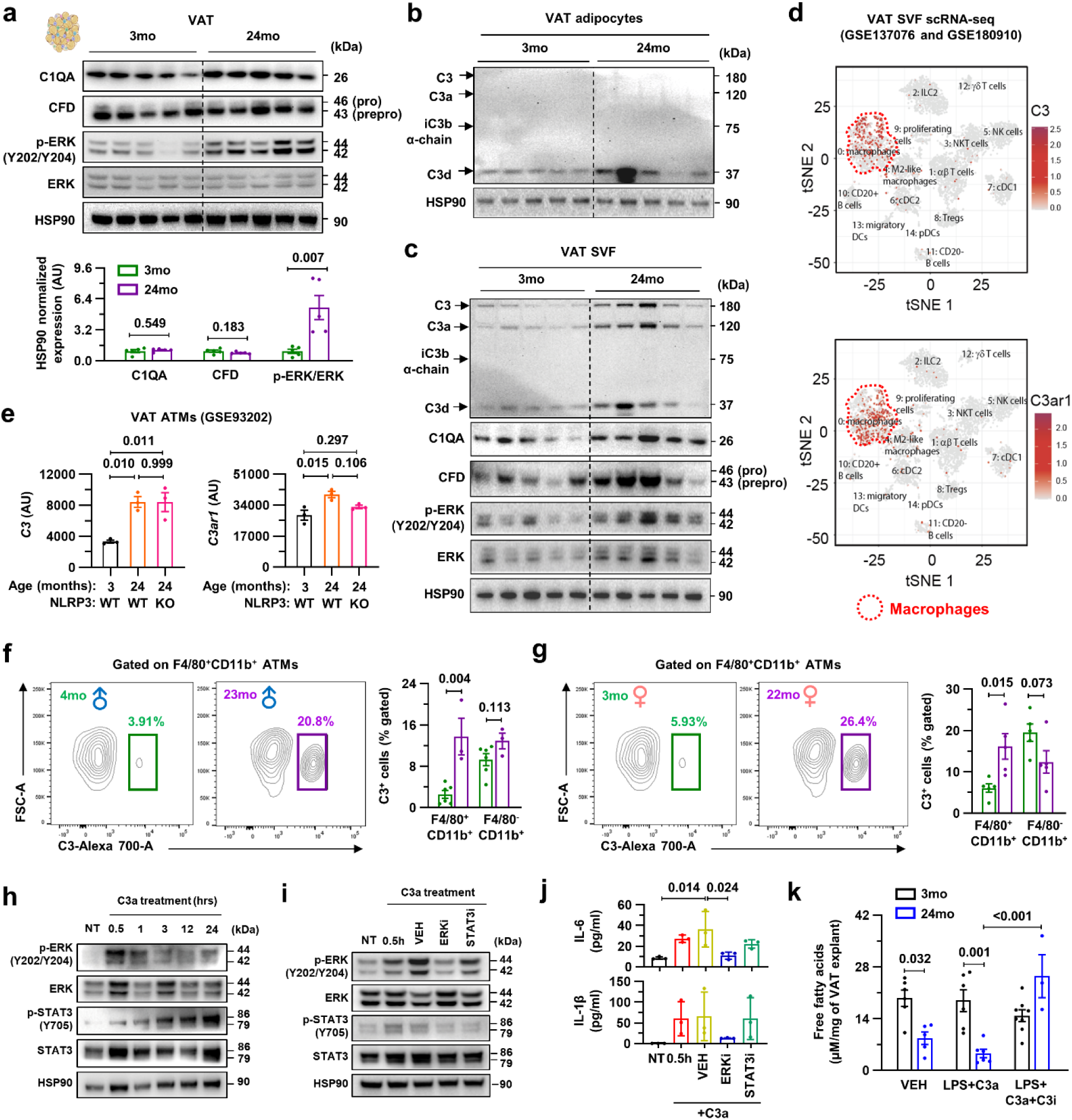
Autocrine C3a signaling in adipose tissue macrophages regulates inflammaging. (a) Representative western blot (top) with densitometry analysis (bottom) of 3- and 24-month-old mice VAT (*n* = 5/group). (b) Representative western blot analysis of 3- and 24-month-old VAT adipocytes (*n* = 5/group). (c) Representative western blot analysis of 3- and 24-month-old VAT stromal vascular fraction (SVF) (*n* = 5/group). (d) Single-cell RNA sequencing analysis of mouse VAT SVF was performed (GSE137076 and GSE180910), and feature plots for *C3* and *C3ar1* expression were presented. (e) Bulk RNA sequencing analyses of VAT ATMs were performed in 3- and 24-month-old wild-type and 24-month-old NLRP3^-/-^ male mice (*n* = 3/group) (GSE93202). *C3* and *C3ar1* expression levels were analyzed. (f,g) Flow cytometry analysis for the C3 expression in F4/80^+^CD11b^+^ VAT ATMs and F4/80^-^CD11b^+^ non-macrophage fraction of 3- and 24-month-old male (f; *n* = 6 for 4-month-old and *n* = 3 for 23-month-old) and female (g; *n* = 5 for 3-month-old and *n* = 5 for 22-month-old) mice with representative panels. (h) Representative western blot analysis of mouse bone marrow-derived macrophages (BMDMs) treated with recombinant C3a (20 ng/ml) for the indicated time points. (i,j) BMDMs were pre-treated with vehicle (DMSO), ERK inhibitor (U0126; 2.5 µM), or STAT3 inhibitor (S3I-201; 25 µM) 1 hour before C3a (20 ng/ml), and subjected to western blot analysis (i) or ELISA (j; *n* = 3/group). (k) VAT explants from young (3-month-old) and old (24-month-old) mice were treated with either vehicle (VEH) or LPS (500 ng/ml) and C3a (40 ng/ml) for 4 hours, and free fatty acid levels were measured in the culture supernatant (*n* = 6 for young VEH, *n* = 6 for young LPS+C3a, *n* = 8 for young LPS+C3a+C3i, *n* = 5 for old VEH, *n* = 6 for old LPS+C3a, and *n* = 3 for old LPS+C3a+C3i). Unpaired two-tailed t-tests (a, f, and g) or One-way ANOVA with multiple comparisons (e, j, and k) were performed, and exact p-values were presented. Data presented as mean±SEM.

To gain further insight into the cellular source of C3, we analyzed our previously reported single-cell RNA sequencing (scRNA-seq) data of VAT resident immune cells^55^. Interestingly, we found that both *C3* and *C3ar1* expressions were enriched in adipose tissue macrophages (ATMs) among the 15 different resident immune cell types (Fig. 5d). In addition, when we analyzed age-associated changes in the expression levels of these genes in ATMs through RNA-seq^56^, which has greater sensitivity than scRNA-seq, both *C3* and *C3ar1* were significantly increased during aging independent of NLRP3 inflammasome (Fig. 5e), suggesting autocrine activation of C3 in ATMs during aging. To confirm the age-associated increase in C3 expression in ATMs at the protein level, we performed flow cytometry analysis (Extended Data Fig. 5c). We observed that more F4/80^+^CD11b^+^ ATMs in aged mice expressed C3 compared to young mice of both sexes, whereas cells in the F4/80^-^CD11b^+^ non-macrophage fraction did not exhibit any difference (Fig. 5f,g).

After confirming that ATMs are key contributors to the VAT-specific age-associated increase in C3a production, we sought to further understand the signaling mechanisms and physiological functions of observed changes. To model this, we utilized bone marrow-derived macrophages (BMDMs) from young male mice. We found that recombinant C3a treatment induced early ERK activation and time-dependent signal transducer and activator of transcription 3 (STAT3) activation in BMDMs, suggesting that BMDMs possess the machinery to react to exogenous C3a (Fig. 5h). In addition, we observed that STAT3 activation by C3a treatment was ERK-dependent, as pre-treatment with an ERK inhibitor also suppressed STAT3 activation in BMDMs (Fig. 5i). In contrast, pre-treatment with a STAT3 inhibitor did not affect ERK activation (Fig. 5i). Consistent with these results and the previously reported pro-inflammatory function of the C3a signaling pathway in macrophages^57^, we found that C3a treatment induced IL-1β and IL-6 production by BMDMs in an ERK-dependent but STAT3-independent manner (Fig. 5j).

Our previous work had found that ATMs actively modulate lipid metabolism in adipose tissue during aging^56^. To determine whether C3a signaling could also affect this phenotype, we performed *ex vivo* experiments with VAT explants from 3- and 24-month-old mice, treating them with C3a in addition to lipopolysaccharide (LPS) to mimic *in vivo* inflamed environments. We found that C3a and LPS treatment did not affect lipolysis in VAT explants from either 3- or 24-month-old mice, although reduced lipolysis was observed in VAT explants from 24-month-old mice compared to 3-month-old mice (Fig. 5k). However, C3 inhibition increased lipolysis in VAT of 24-month-old mice (Fig. 5k), suggesting the potential role of complement pathways in improving metabolic pathways during aging. Collectively, these data may indicate that C3a is produced by ATMs and promotes age-associated inflammation in an autocrine manner.

### C3 inhibition improves healthspan and reduces inflammaging

The above findings guided us to investigate whether C3 inhibition could mimic the immunometabolic benefits of CR *in vivo*. For this purpose, we intraperitoneally administered either the vehicle or 1 mg/kg of the C3 inhibitor, AMY-101, a small peptide compound that has recently concluded a phase IIa clinical trial for patients with periodontal inflammation^58,59^. This was first conducted on young (3-month-old) male mice weekly for 6 weeks to assess any potential side effects of the inhibitor.

We found that body weight was well maintained throughout the experiments, suggesting that the dosage of the C3 inhibitor used in this study was safe and well-tolerated (Extended Data Fig. 6a,b). In addition, when we measured fat mass, lean mass, and weights of peripheral tissues, including the SAT, VAT, liver, spleen, and kidney, we did not observe significant differences between the vehicle and C3 inhibitor-treated groups (Extended Data Fig. 6c-e). Similarly, there was no significant difference in the glucose tolerance test between the two groups (Extended Data Fig. 6f). To define the immunological effects of C3 inhibitor treatment in healthy young mice, we quantified various cytokines and chemokines in the serum. Although macrophage inflammatory protein (MIP)-1β showed a modest increase in C3 inhibitor-treated compared to vehicle-treated mice, the serum levels of IL-1β, IL-6, TNF-α, monocyte chemotactic protein (MCP)-1, and granulocyte-macrophage colony-stimulating factor (GM-CSF) were comparable between the two groups (Extended Data Fig. 6g-l). In addition, the expression levels of the NLRP3 inflammasome-associated genes, such as *Nlrp3, Casp1, Il1b*, and *Il18*, were not affected by C3 inhibitor treatment in the VAT of adult mice (Extended Data Fig. 6m-p). These results indicate that the C3 inhibitor does not provoke any unexpected immune or metabolic disruptions in young mice.

Based on this data, we next examined the effects of the C3 inhibitor on aged (20-month-old) male mice, which harbor elevated C3a expression in the VAT ATMs and immune-metabolic dysfunctions. Although there was a tendency towards lower body weight and fat mass in the C3 inhibitor-treated group compared to the vehicle-treated group, we did not find significant differences in body weight, fat mass, or lean mass between the groups (Fig. 6a-d). Consistent with this and the ex vivo experiments with the VAT explants, we did not observe significant differences in the phosphorylation of hormone-sensitive lipase (HSL) and adipose triglyceride lipase (ATGL) levels (Fig. 6e). However, the activation of ERK signaling, downstream of C3AR1, was significantly mitigated in the VAT by C3 inhibitor treatment in aged mice (Fig. 6e). Interestingly, the C3 inhibitor treatment significantly improved healthspan, including glucose tolerance and grip strength, and there was a tendency to enhance motor coordination in the rotarod test (Fig. 6f-h).

**Fig. 6.**
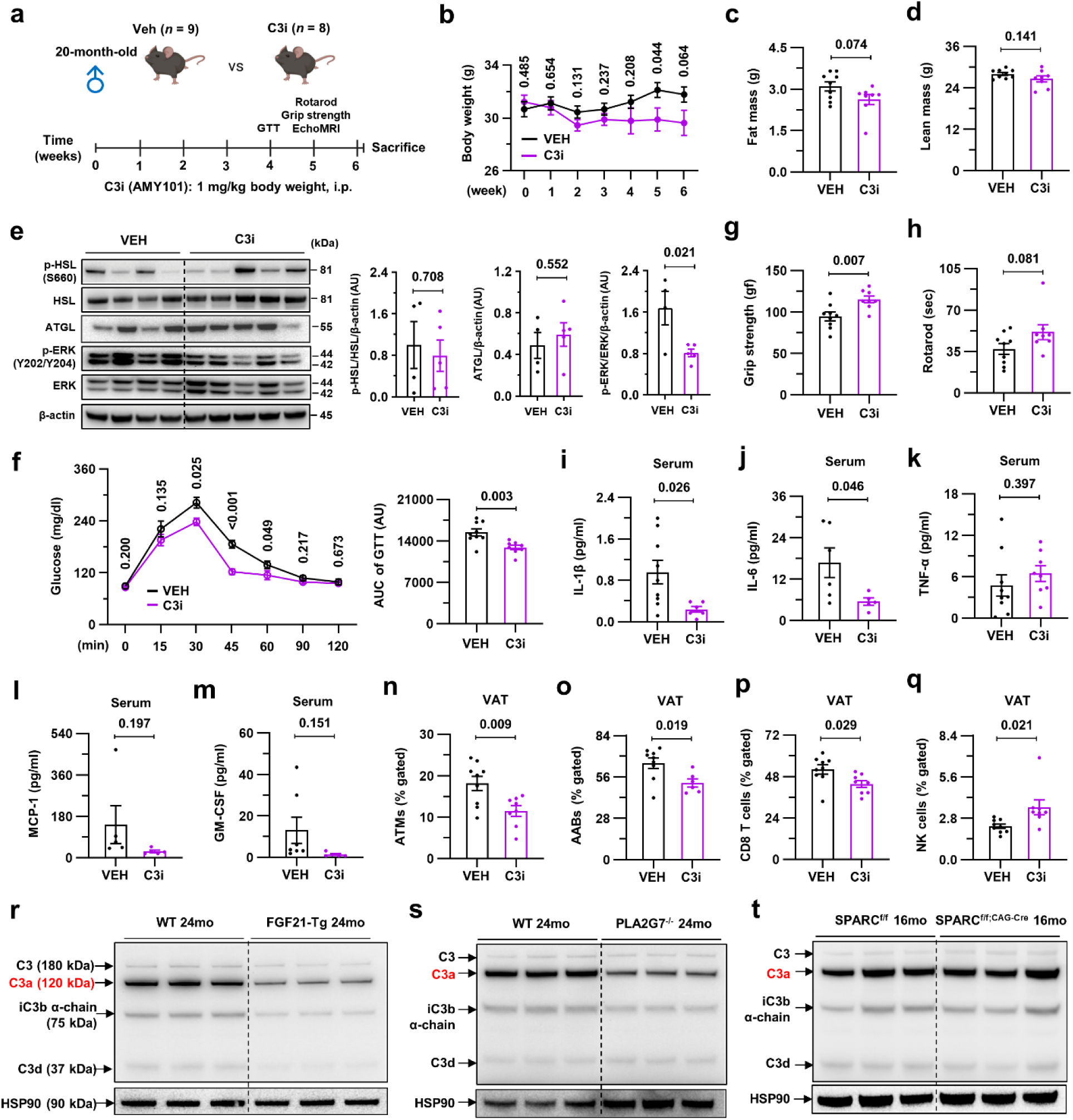
C3 inhibition improves healthspan and reduces inflammaging. (a-q) Aged (20-month-old) male mice were randomly allocated and intraperitoneally injected with either vehicle or the C3 inhibitor (AMY-101; 1 mg/kg) weekly for 6 weeks and analyzed (*n* = 9 for vehicle and *n* = 8 for C3 inhibitor). (a) Schematic diagram for the experimental design. (b) Body weight measurement. (c,d) Fat mass (c) and lean mass (d) were measured by EchoMRI. (e) Representative western blots with densitometry (*n* = 4 for vehicle and *n* = 5 for C3 inhibitor). (f) A glucose tolerance test was performed. (g,h) Healthspan was analyzed through grip strength (g) and the rotarod test (h). (i-m) Serum levels of IL-1β (i), IL-6 (j), TNF-α (k), MCP-1 (l), and GM-CSF (m) were analyzed by Luminex assay. Only detected values were presented. (n-q) Flow cytometry analysis of VAT SVF. The frequencies of ATMs (n), AABs (o), CD8 T (p), and NK cells (q) were presented. (r-t) Representative C3 western blot analysis of the VAT of 24-month-old WT and FGF21-Tg (r), 24-month-old WT and PLA2G7^-/-^ (s), and 16-month-old SPARC^f/f^ and SPARC^f/f;CAG-Cre^ (t) mice (*n* = 3/group). Unpaired two-tailed t-tests were performed, and exact p-values were presented. Data presented as mean±SEM.

We next analyzed the impact of C3 inhibition on immunological hallmarks of aging. Among the immune effectors, we observed significantly reduced serum levels of IL-1β and IL-6, key drivers of inflammaging^60^, in the C3 inhibitor-treated compared to vehicle-treated group, whereas TNF-α, MCP-1, and GM-CSF levels were comparable (Fig. 6i-m). Given that VAT ATMs were the major source of age-associated C3 elevation, we next investigated the effects of C3 inhibition on multiple immune cell types in the VAT utilizing multi-parametric flow cytometry (Extended Data Fig. 7a). We found that pharmacological inhibition of C3 in aged mice suppressed the frequencies of ATMs, adipose aged B cells (AABs), and CD8 T cells, all of which have been implicated in age-associated inflammation or complement activation (Fig. 6n-p)^56,61–63^. In contrast, the frequencies of natural killer (NK) cells were higher in the C3 inhibitor-treated mice compared to the vehicle group, while the frequencies of Ly6C^+^ monocytes, eosinophils, γδ T cells, and NKT cells remained unchanged (Fig. 6q and Extended Data Fig. 7b-e). These results indicate that the inhibition of C3 restrains key drivers of inflammaging.

Given that overexpression of FGF21 and genetic inhibition of PLA2G7 or secreted protein acidic and cysteine-rich (SPARC) have been established as interventions to extend healthspan or potentially lifespan by restraining inflammaging^5,6,64–66^, we investigated C3a in the VAT of aged mice overexpressing FGF21 and lacking PLA2G7 and SPARC to determine whether the reduction in C3a levels is a common feature of healthy longevity. Of note, 24-month-old FGF21 transgenic (FGF21-Tg) and PLA2G7 knockout mice exhibited significantly reduced C3 cleavage compared to age-matched controls (Fig. 6r,s and Extended Data Fig. 7f,g). However, 16-month-old SPARC knockout mice showed comparable expression of C3 and its cleaved forms to control mice (Fig. 6t and Extended Data Fig. 7h), suggesting that the pro-longevity effects of FGF21 overexpression and PLA2G7 reduction may involve a reduction in C3 levels. In contrast, the SAT from 18-month-old adipocyte-specific FGF21-overexpressed (ADN-iFGF21-Tg) mice did not show any reduction in C3 levels (Extended Data Fig. 7i), highlighting that C3 activation is specific to the VAT. Collectively, these data suggest that the inhibition of C3 extends healthspan by regulation of immunometabolic mechanisms in aged mice.

## Discussion

CR, the most effective non-pharmacological pro-longevity intervention, has been shown to improve healthspan and extend lifespan in many species, including rodents and primates^5,6,15^. In the present series of studies, including unbiased proteomics analysis of a human CR clinical trial, we propose that targeting C3 activation could be a CR mimetic that enhances the healthspan and potentially lifespan (Extended Data Fig. 8). Under homeostatic conditions, inflammatory cues result in complement activation and resolve the damage. However, elevated C3 levels during aging are detrimental to longevity, and a reduction in C3 levels has been shown to improve age-related kidney changes in mice and extend lifespan in *C. elegans*^17,67^. Therefore, increased age-associated complement activation can potentially be targeted to health outcomes in older adults.

Aging is associated with increased visceral adiposity, a hallmark of aging, characterized by homeostatic perturbations, weakened stress resilience, metabolic decline, and chronic inflammation^68^. Among the primary goals of translational geroscience research is to delineate mechanistic insights of aging and to discover or target the pathways with interventions that mitigate functional decline in aging. To achieve this goal, recent discoveries have established genetically manipulated FGF21 and PLA2G7 mouse models that exhibit extended healthspan^5,69,70^, and finding the common mechanistic targets across these established interventions could accelerate progress in geroscience. Our studies, based on human CR intervention data, identified that adipose tissue complement deactivation is a common inflammaging checkpoint shared among non-pharmacological and genetic interventions. Given that many complement inhibitors are clinically being tested and complement activation has been implicated in multiple aging-related diseases, including Alzheimer’s disease, age-related macular degeneration, and osteoarthritis^71^, pharmacological inhibition of C3 activation could be a feasible approach to delay systemic physiological and functional declines during aging.

Indeed, in line with reduced complement proteins in the CALERIE participants, we found that pharmacological inhibition of C3 improved the metabolic fitness, motor activity, and age-related inflammation in aged mice. However, one potential pitfall in the long-term clinical use of C3 inhibitors is the existence of non-canonical intracellular C3 signaling pathways, which regulate essential cellular processes, including energy metabolism, lipid metabolism, cell survival, and efferocytosis^51^. Although we did not observe any potential side effects in our in vivo experiments in both young and aged mice, there was a tendency toward impaired glucose tolerance and inflammation in young mice treated with the C3 inhibitor. Therefore, further studies are required to determine the most efficient and safe dose of the C3 inhibitor for long-term clinical use.

Of note, our data implicate the VAT as a major contributor to age-associated increased complement activation-mediated inflammation. In addition, we also report that ATMs are central in displaying age-associated complement cleavage by utilizing ERK-dependent STAT3 activation. However, how macrophage-specific C3 regulates the adipose tissue microenvironment during the aging process requires further investigations with genetically modified mouse models. Since complement proteins are involved in multiple physiological events and C3AR1 is present in many cell types, whether macrophage-specific C3 effector functions rely more on autocrine, paracrine, or endocrine signaling during the aging process remains to be answered.

In summary, our reverse translational approach demonstrates that deactivation of complement serves as an inflammaging checkpoint and highlights the pivotal role of maintaining adipose tissue homeostasis during aging. Further mechanistic studies will provide more insights into targeting specific downstream pathways of C3 and lead to the development of pharmacological interventions to extend healthspan and lifespan.

## Methods

### Human participants

The samples used for the plasma proteomics were from the participants of the CALERIE Phase II study, a multicenter and randomized controlled trial. Plasma samples were collected at the baseline and after 2 years of 14% caloric restriction (CR) from a total of 42 middle-aged non-obese healthy individuals. All the detailed characteristics of the participants were listed in Extended Data Table 1. Serum samples of a total of 17 young (21 to 30 years old) and 23 older adults (>65 years old) were collected at Yale Hospital to quantify the C3. All clinical studies were performed under the approved guidelines by the Institutional Review Board at Pennington or Yale, with informed consent from participants.

### Mice

Young (3 to 5 months old) and aged (20 to 24 months old) WT male and female C57BL/6J mice were purchased from the Jackson Laboratory, and C57BL/6N mice were provided by the National Institute on Aging (NIA) Aged Rodent Colony. All mice used in this study were maintained in a ventilated cage rack in specific pathogen-free conditions at Yale School of Medicine. Mice were group housed and fed with a standard vivarium chow diet and water *ad libitum* and maintained under a regular 12-hour light/dark cycle. C57BL/6N mice were mainly used for the mouse experiments, otherwise stated in the figure. The sex of the mice used was represented in each figure and legend. For C3 inhibitor treatment *in vivo* experiments, 3-month-old and 20-month-old C57BL/6N male mice were randomly allocated to vehicle or C3 inhibitor (AMY-101; 1 mg/kg) (GLPBIO) group, and the treatment was intraperitoneally administered weekly for 6 weeks. Mice were sacrificed a week after the last inhibitor treatment. The Yale Institutional Animal Care and Use Committee reviewed and approved all animal procedures.

### SomaScan 7K assay

Plasma samples of CALERIE participants (*n* = 42) were subjected to the SomaScan 7K assay. The initial assay and normalization followed SomaLogic’s pipeline. Briefly, SomaScan assay data are initially normalized using hybridization controls to reduce variation within the run caused by readout steps. Subsequently, median signal normalization is performed across pooled calibrator replicates within the run to address within-run technical variations in the calibrator signals before employing them for scaling calculations. The ratios of the reference value to the median of replicates for each SOMAmer are calculated and decomposed into two components: plate scale – representing the median ratio, and calibration scale – representing the recalculated set of scale factors specific to each SOMAmer reagent. The plate scale adjusts for overall signal intensity differences between runs, while calibration adjusts for reagent-specific assay differences between runs. Median signal normalization is conducted using Adaptive Normalization by Maximum Likelihood (ANML) for plasma.

For proteomic expression, we used ANML normalization values provided by SomaLogic. The values were log2-transformed, and proteins that failed internal SomaScan quality control were removed. All the SOMAmers that failed quality control were listed in Extended Data Table 2. To identify proteins differentially expressed under CR, we fitted linear models using the limma R package (version 3.56.2)^72^, including donor as a blocking factor for paired testing.

For hallmarks of aging analysis, the list of genes associated with each aging hallmark pathway was obtained from the Human Aging Atlas^35^. Heatmaps were generated using the z-scored log2 fold change of ANML normalized values. Scoring analysis was conducted by calculating the average value of all the genes involved in each specific pathway. Statistical significance was assessed using a two-tailed paired t-test, and adjusted p-values were used. The proteomic age of each organ was estimated using the ‘organage’ package^36^. Input data included SOMAmer ID and ANML normalized values in RFU units provided by SomaLogic. The calculated z-scored age gaps from the ‘organage’ package were used to predict whether the aging of each organ was accelerated or delayed following caloric restriction.

Pathway enrichment of the resulting protein signatures was performed with the fgsea R package (version 1.26.0)^73^, using the limma t-statistic as the ranking metric. Volcano plots were generated with ggplot2 (version 3.5.1). For PCA, expression values were first normalized via variance stabilizing normalization using the vsn R package (version 3.68.0)^74^, after which PCA was performed with the base R prcomp function and visualized in ggplot2 (version 3.5.1). R version 4.3.0 was used for all the analyses.

### Ingenuity Pathway Analysis (IPA)

The fold change and p-values from the SomaScan 7K assay were used as inputs for IPA analysis. The upstream regulator analysis for NF-κB and the canonical pathway analysis for the ‘complement system’ were conducted according to the manufacturer’s instructions.

### ELISA and Luminex

The serum protein levels of C3 (Abcam) in humans and mice were assessed using ELISA in both sexes, male and female. Serum C3a level (Thermo) was measured in male and female mice only. For the Luminex assay, 25 μL of collected serum was used to detect IL-1β, TNFα, IL-6, MCP-1, MIP-1β, and GM-CSF by the mouse custom ProcartaPlex6-plex kit (Thermo Fisher Scientific) on the Luminex 200 analyzer.

### Immunoblot

Western blots were carried out on the whole cell lysate (WCL) from liver, kidney, spleen, subcutaneous adipose tissue (SAT), brown adipose tissue (BAT), and visceral adipose tissue (VAT) collected from young and old mice of both sexes, which were homogenized in RIPA buffer containing phosphatase and protease inhibitor cocktail. Complement C3 (ab200999; Abcam), C1QA (ab155052; Abcam), Factor-D/Adhipsin (MAB5430-SP; R&D), p-STAT3 (9145; Cell Signaling), total STAT3 (4904; Cell Signaling), p-Erk (4377S, Cell Signaling), total Erk (4695S, Cell Signaling), HSP-90 (60318-1-Ig, Proteintech), β-actin (4967L; Cell Signaling) Phosphorylated HSL S536 (Cell Signaling; 4139S), Phosphorylated HSL S660 (Cell Signaling; 45804S), Total HSL (Cell Signaling; 4107S) as primary antibodies and HRP-conjugated goat anti-rabbit or anti-mouse as secondary antibodies were used to detect proteins using ChemiDoc MP Imaging System (Bio-Rad).

### Adipose tissue digestion and flow cytometry analysis

VAT was digested, and SVF cells were stained as described^6^. Cells were first incubated with ViaKrome 808 Fixable Viability Dye (catalog no. C36628, BECKMAN COULTER) and further stained with following surface antibodies-Fc receptor (CD16/32) blocking (14-0161-86, Invitrogen) for 10 minutes, CD45-BV421 (Clone 30-F11, catalog no. 563890), CD11c-BV750 (Clone N418 catalog no. 117357), B220-BV711 (Clone RA3-6B2 catalog no. 103255), Ly6C-AF700 (Clone HK1.4 catalog no. 128024), PerCP/Cyanine5.5-TCR β chain (Clone H57-597 catalog no. 109228), PerCP/Cyanine5.5-CD45R/B220 (Clone RA3-6B2 catalog no. 103236) (all from BioLegend); Siglec-F-BV605 (Clone E50-2440 catalog no. 740388), CD11b-BUV395 (Clone M1/70 catalog no. 563553), CD3-PE-Cy7 (Clone 17A2 catalog no. 560591), γδTCR-BV650 (Clone GL3 catalog no. 563993), CD8a-BUV615 (Clone 53-6.7 catalog no. 613004), (all from BD Biosciences); F4/80-StarBrightViolet570 (Clone CI:A3-1 catalog no. MCA497) (from Bio-Rad) and CD45-NovaFluor Blue 660-120S (Clone 30-F11 catalog no. M005T02B08), NK1.1-PE (Clone PK136 catalog no. 2-5941-83) (from Thermo Fisher Scientific). When required, C3-AF700 (Clone 11H9 catalog no. NB200-540AF700) antibody was used for intracellular staining (from Novus Biologicals). Samples were acquired on a Becton Dickinson Symphony instrument, and data were analyzed using FlowJo software (Treestar).

### Ex vivo lipolysis

Young and old VAT explants (around 15 mg) were collected in a 96-well plate and cultured in lipolysis buffer with 500 ng/ml of LPS & 40 ng/ml of recombinant C3a for 4 hours, followed by 2 hours of C3i treatment (10 uM) at 37^°^C. Free fatty acid and glycerol assay (both from Sigma) was performed according to the manufacturer’s instructions.

### BMDM culture

BMDMs were prepared from male mice between 8-12 weeks of age as described^6^. BMDMs were treated with LPS (500 ng/ml) before recombinant C3a protein (20 ng/ml) treatment. For blocking ERK or STAT3 activation, BMDMs were treated with ERK inhibitor U0126 (2.5 uM) or STAT3 inhibitor S3I-201 (25 uM) for 1 h before LPS or C3a stimulation.

### Glucose tolerance test (GTT)

Following a 16-hour fast, young mice (1g/kg/bw) and aged mice (0.4g/kg/bw) received intraperitoneal injections of a 25% and 10% glucose solution (Sigma), respectively. Mice were subjected to this GTT (Contour Next blood glucose strip and meter, Diabetic Corner) test 3 days after receiving the C3 inhibitor injection during the 4^th^ week.

### RNA Extraction, cDNA Generation, and RT-qPCR

Total RNA was extracted utilizing Trizol and subsequently purified with the RNeasy Kit (Qiagen) in accordance with the manufacturer’s guidelines. cDNA synthesis was performed utilizing the iScript cDNA Synthesis Kit (BioRad) in accordance with the manufacturer’s guidelines. 1-2 µg of cDNA was diluted to 5 ng/mL and amplified using specified primers in a 20 µL reaction with Power SYBR Green PCR Master Mix (Applied Biosystems). Gene expression analysis was conducted using a LightCycler 480 II instrument (Roche). For each gene, mRNA expression was calculated relative to *Actb* and *Hprt1* expression. Primers used in this study are listed in Extended Data Table 3.

### EchoMRI, grip strength, and rotarod test

Mice were subjected to EchoMRI to measure body composition (fat mass, lean mass, and free water) and grip strength/rotarod test to conduct behavioral testing as described^6^. The TSE system’s grip strength meter was used to conduct the grip strength test. When mice were holding a bar in the meter, their forelimbs’ maximum strength (gram force, or gf) was measured three times, and the average of them was used to calculate the strength. A rotarod test was conducted using an accurotor rotarod (AccuScan Instruments). The average of the three tests that were conducted one day following training to determine how long the mice could remain on the rotarod was used for analysis. 400 (x 0.1 RPM) was the maximum speed, and 300 (sec) was the time to maximum speed.

### Free fatty acid (FFA) and glycerol measurements

Blood was collected through cardiac puncture at the time of sacrifice and incubated at room temperature for 2 h, followed by centrifugation at 3,000 rpm for 20 min at 4 ^°^C to collect the serum. Serum FFA and glycerol levels were measured using Free Fatty Acid Assay Kit (MAK466; Sigma) and Glycerol Assay kit (MAK117; Sigma), respectively, per manufacturer’s instructions.

### Quantification and statistical analysis

The exact numbers of animals and statistical methods used in each experiment can be found in the figure legends. For the plasma proteomics analysis, adjusted p-values were used. For the in vitro and in vivo experiments, a two-tailed Student t-test (between the two groups) or One-way ANOVA with multiple comparisons (more than three groups) was performed. All statistical tests were performed using GraphPad Prism 9 for Windows (GraphPad Software, San Diego, United States of America). The exact p-values were presented in each figure. Data presented as mean±SEM.

### Data Availability

Previously published datasets that were used in this study are indicated by the GEO number in the figures.

## Acknowledgements

We thank all investigators and staff who coordinated and executed the CALERIE-II clinical trial. The research in the Dixit Lab was supported in part by NIH grants AG031797, AG045712, P01AG051459, AR070811, and Cure Alzheimer’s Fund (CAF).

## Author Contributions

M.M. conducted western blot analysis, *in vitro* experiments with macrophages, *in vivo* experiments with mice, prepared figures, and wrote the first draft of the manuscript. H.H.K. analyzed the human plasma proteomics data, prepared and arranged figures, and wrote the first draft of the manuscript. Y.Y. and T.D. prepared the materials for the proteomics analyses and organized the clinical information of participants. E.G.H. assisted *in vivo* experiments with young mice. K.Z. and M.N.A. performed the initial analysis of the human plasma proteomics. I.S. and M.N.A. analyzed the single-cell RNA sequencing analysis of the VAT SVF. C.G. and P.E.S. participated in adipose complement experiments, including providing transgenic mice and tissue samples. S.M. and A.C.S. collected and provided human samples for C3 ELISA. E.R. was involved in the design and execution of the CALERIE-II trial and was the PI of the Pennington Biomedical site of CALERIE. V.D.D. conceived the project, helped with data interpretation, and wrote the manuscript.

