## Supplementary figures and tables for "Exoproteome of calorie-restricted humans identifies complement deactivation as an immunometabolic checkpoint reducing inflammaging"

### EXTENDED DATA FIGURES

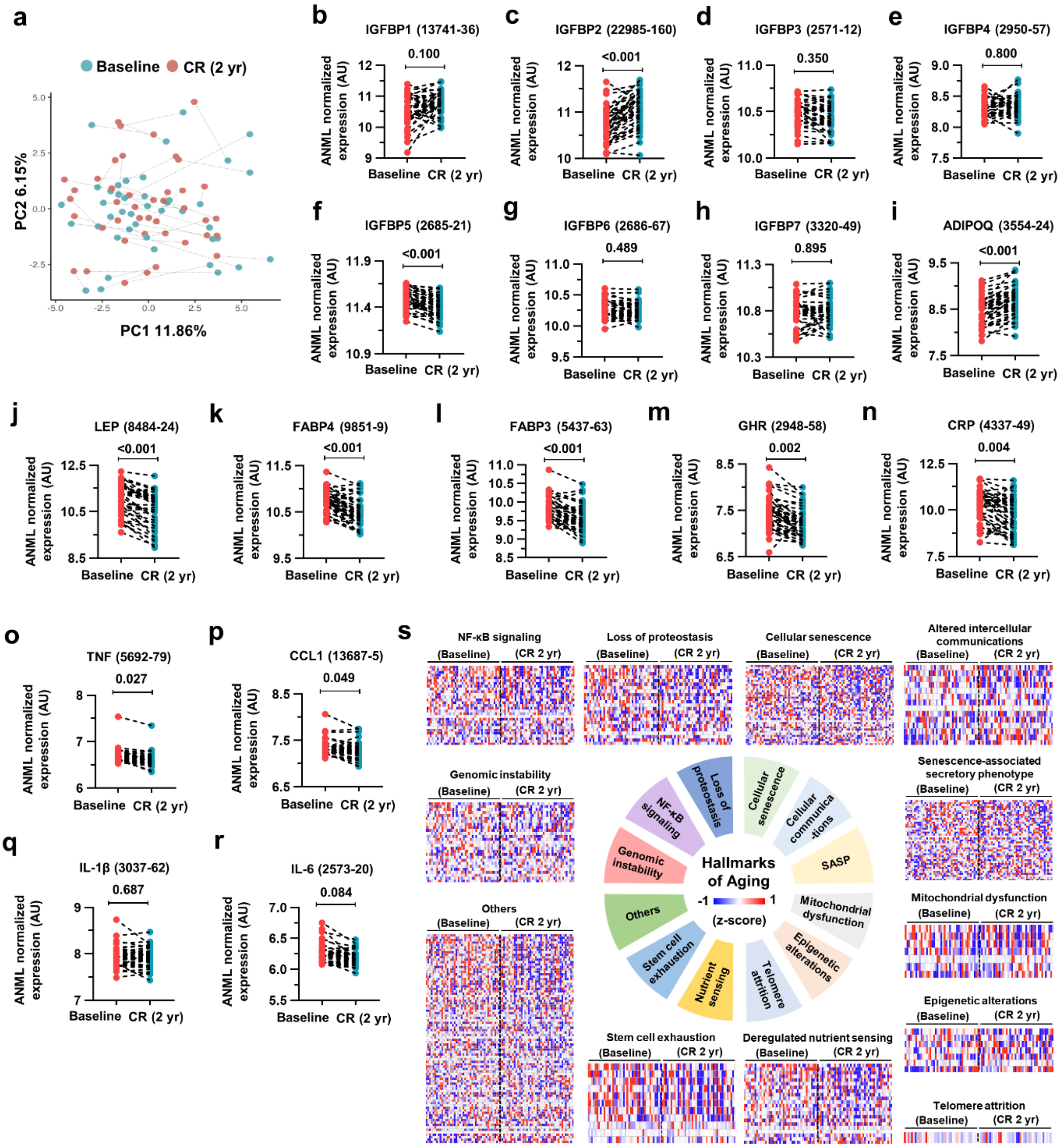


**Extended Data Fig. 1. CR in healthy individuals increases IGFBP2 and adiponectin but reduces leptin, FABPs, and inflammatory molecules in plasma.**

(a-s) SomaScan 7K proteomics was performed in the plasma of healthy individuals at baseline and 2 years after 14% caloric restriction (CR) (*n* = 42). (a) PCA plot. (b-r) The CR-mediated effects on the normalized expression levels of the indicated protein with SOMAmer ID. Paired two-tailed t-tests were performed, and exact adjusted p-values were presented. (s) CR-associated changes in 12 hallmarks of aging pathways were analyzed, and heatmaps for each pathway were presented.

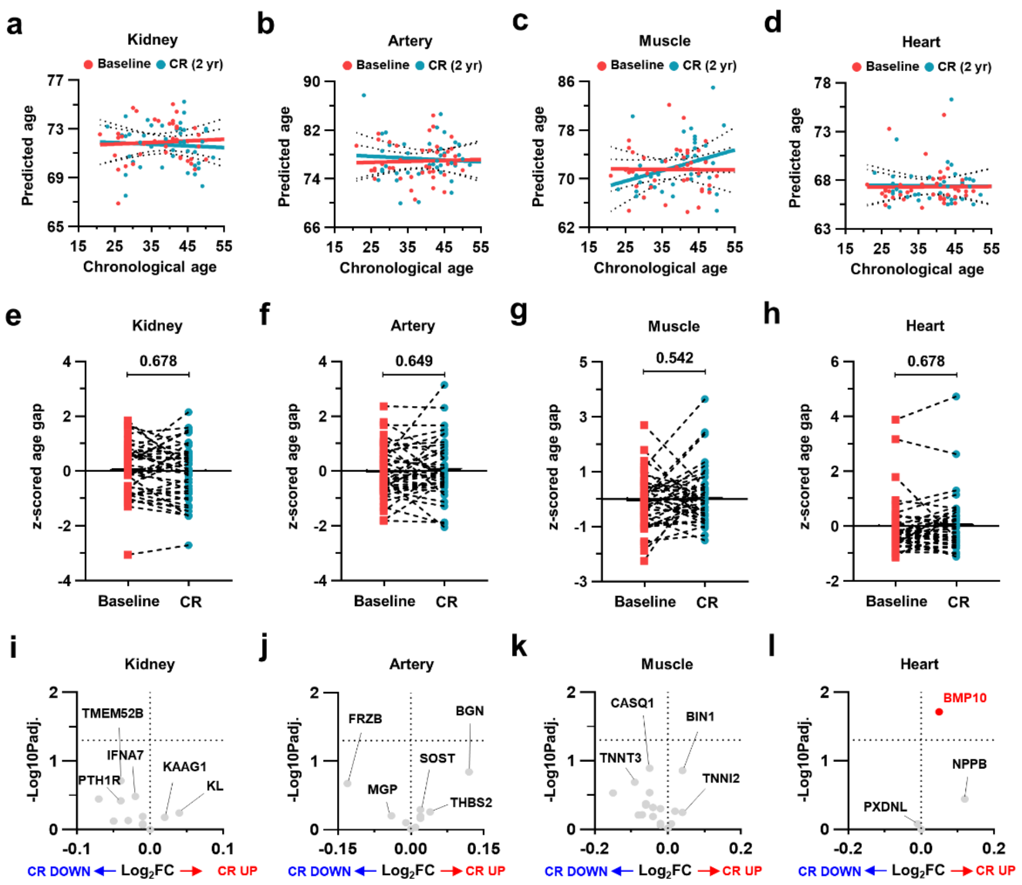


**Extended Data Fig. 2. CR does not affect the proteomic aging of the kidney, artery, muscle, and heart.**

(a-l) Organ-specific aging signatures were analyzed based on the plasma proteome. (a-d) Linear regression analysis was performed to compare the predicted proteomic age before and after CR in the kidney (a), artery (b), muscle (c), and heart (d). (e-h) Organ age gap was analyzed in the kidney (e), lung (f), pancreas (g), and intestine (h). Paired two-tailed t-tests were performed, and exact adjusted p-values were presented. (i-l) Volcano plots for the organ-specific proteins in the kidney (i), artery (j), muscle (k), and heart (l).

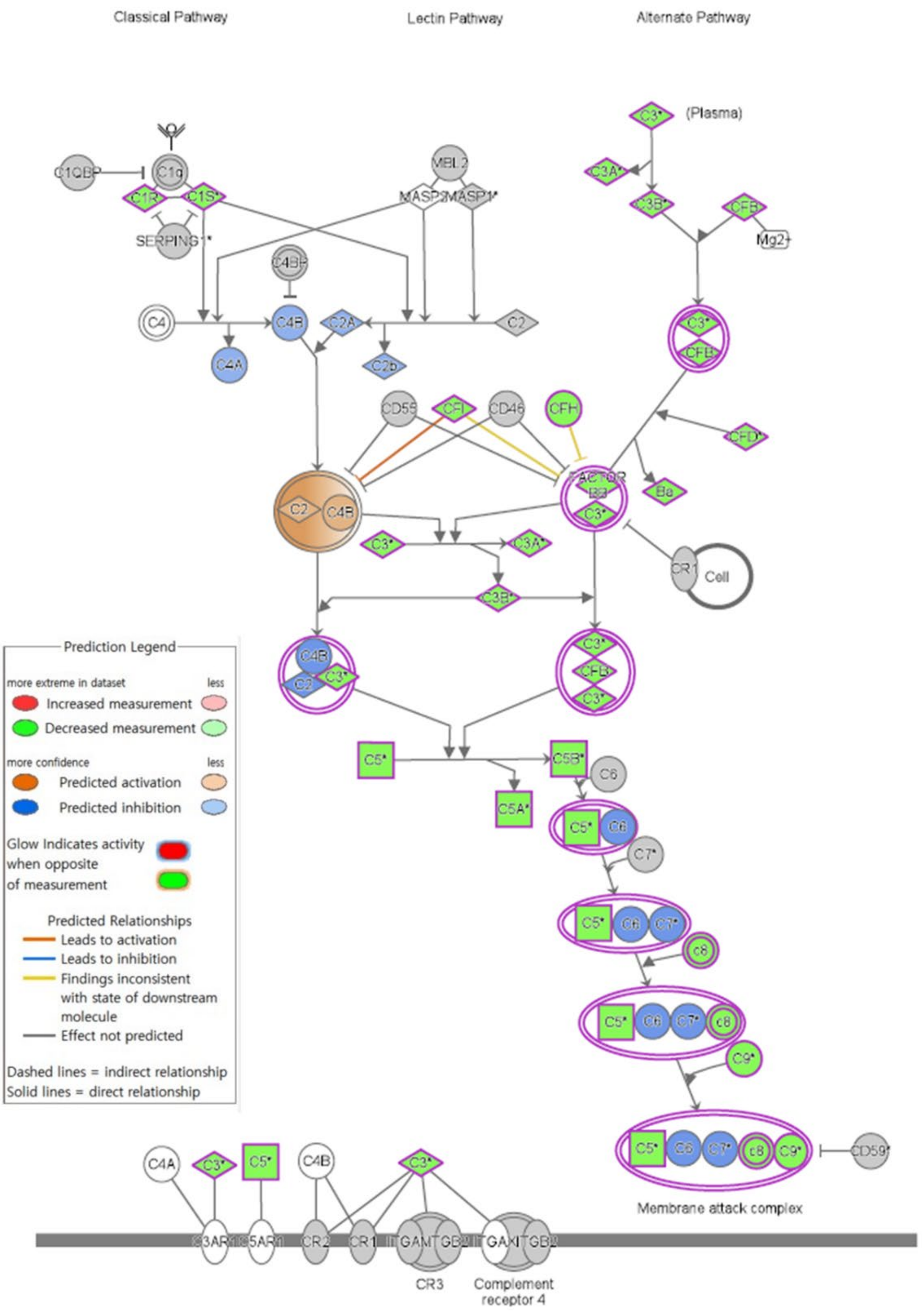


**Extended Data Fig. 3. CR in healthy individuals dampens complement pathways.**

The plasma proteome-based prediction for CR-induced changes in the complement system by Ingenuity Pathway Analysis. Most of the components in the complement system were downregulated by CR.

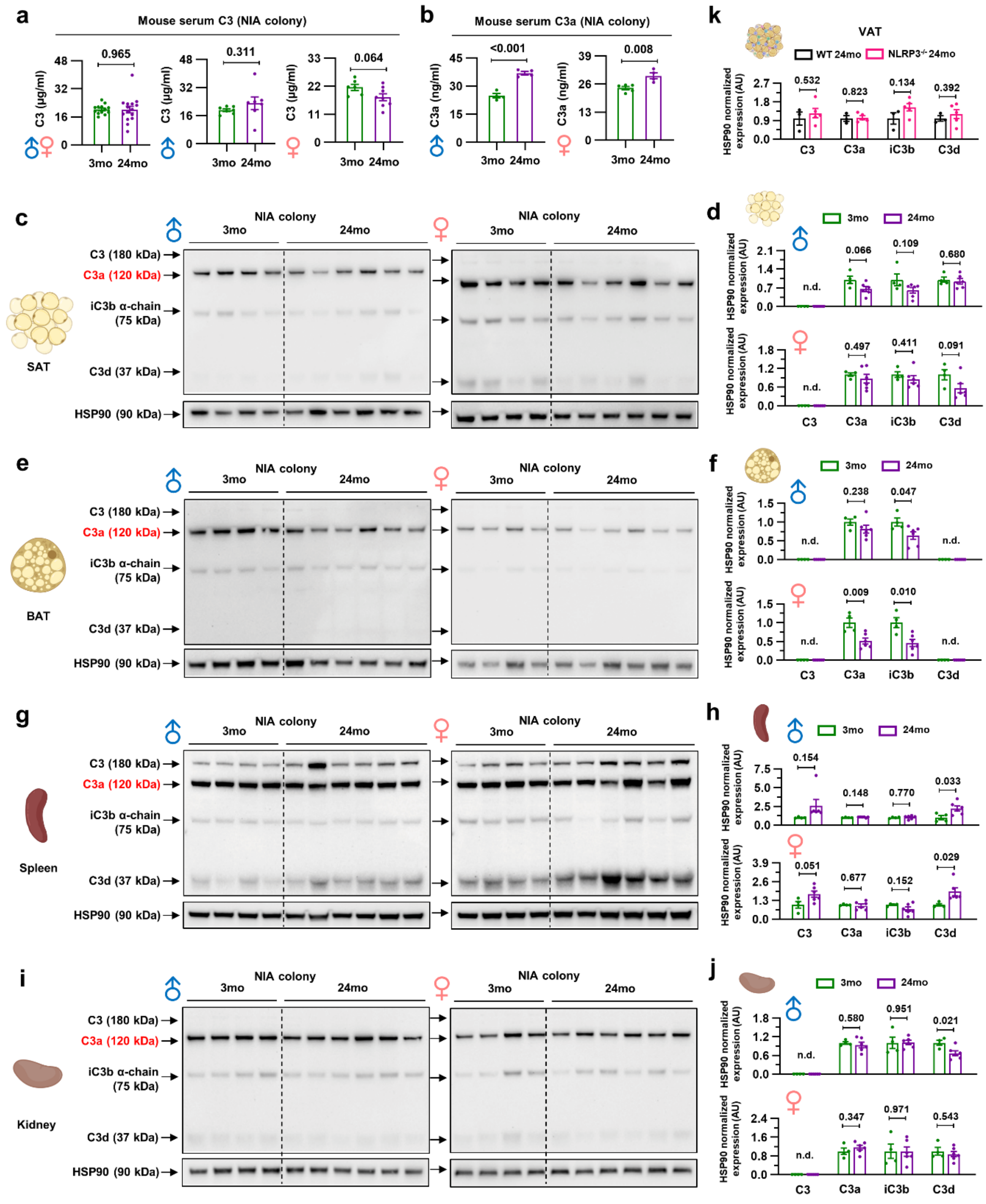


**Extended Data Fig. 4. Age-associated increase in C3a levels was not derived from SAT, BAT, spleen, and kidney in mice.**

(a) Mouse serum samples were analyzed for C3 ELISA in both sexes ($n$ = 13 for 3-month-old and $n$ = 15 for 24-month-old), males ($n$ = 7/group), or females ($n$ = 6 for 3-month-old and $n$ = 8 for 24-month-old). (b) Mouse serum samples were analyzed for C3a ELISA in males ($n$ = 4/group) and females ($n$ = 5 for 3-month-old and $n$ = 4 for 24-month-old). (c-j) Representative

C3 western blot analysis of the various mouse tissues (*n* = 4 for 3-month-old and *n* = 6 for 24-month-old in both sexes). (c,d) C3 western blot analysis of mouse subcutaneous adipose tissues (SAT). (c) Representative blots for C3 in both sexes. (d) Densitometry analysis of male (top) and female (bottom) mice. (e,f) C3 western blot analysis of mouse brown adipose tissues (BAT). (e) Representative blots for C3 in both sexes. (f) Densitometry analysis of male (top) and female (bottom) mice. (g,h) C3 western blot analysis of mouse spleen. (g) Representative blots for C3 in both sexes. (h) Densitometry analysis of male (top) and female (bottom) mice. (i,j) C3 western blot analysis of mouse kidney. (i) Representative blots for C3 in both sexes. (j) Densitometry analysis of male (top) and female (bottom) mice. (k) Densitometry analysis of the VAT of 24-month-old WT and $NLRP3^{-/-}$ mice (*n* = 3/group). Unpaired two-tailed t-tests were performed, and exact p-values were presented. Data presented as mean±SEM.

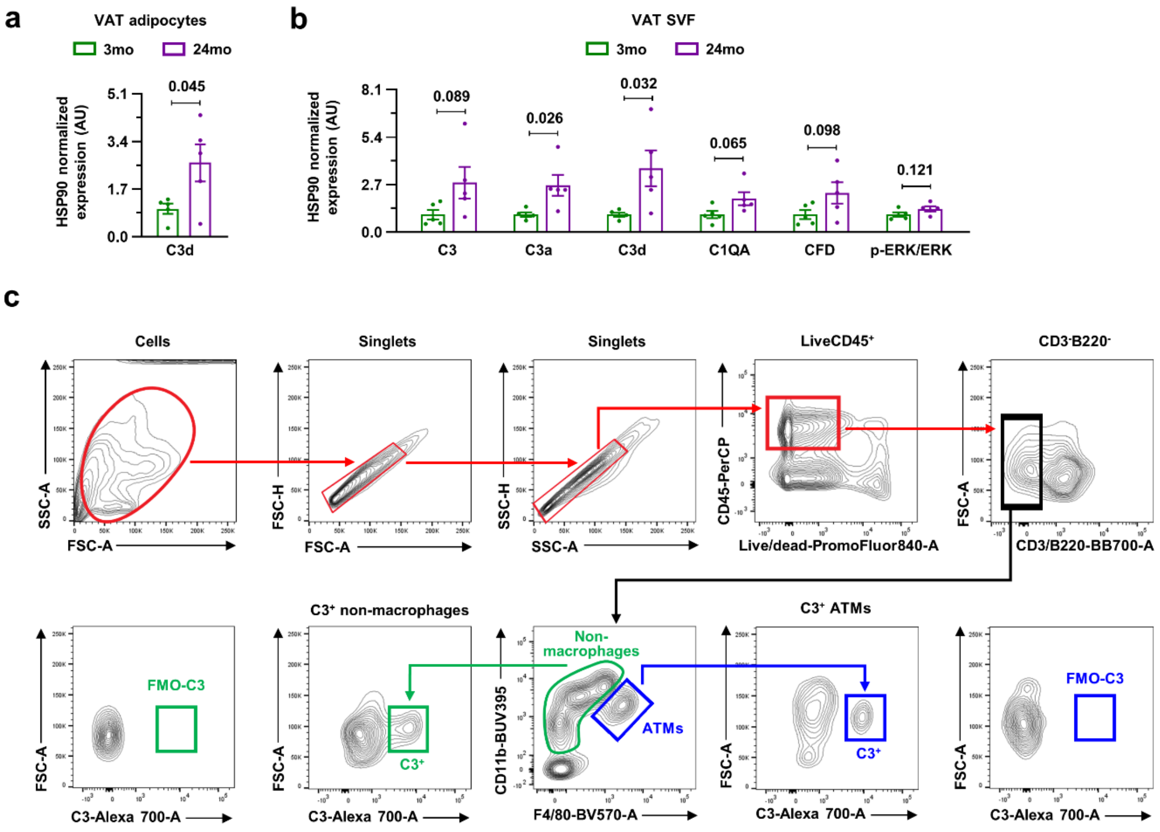


**Extended Data Fig. 5. Age-associated complement activation in the VAT occurs in the non-adipocyte fraction.**

(a) Densitometry analysis of the VAT adipocyte (*n* = 5/group). (b) Densitometry analysis of VAT SVF (*n* = 5/group). (c) Gating strategy for analyzing C3-expressing F4/80$^+$CD11b$^+$ ATMs and F4/80$^-$CD11b$^+$ non-macrophage fractions in the VAT, along with fluorescence minus one (FMO) control for C3.

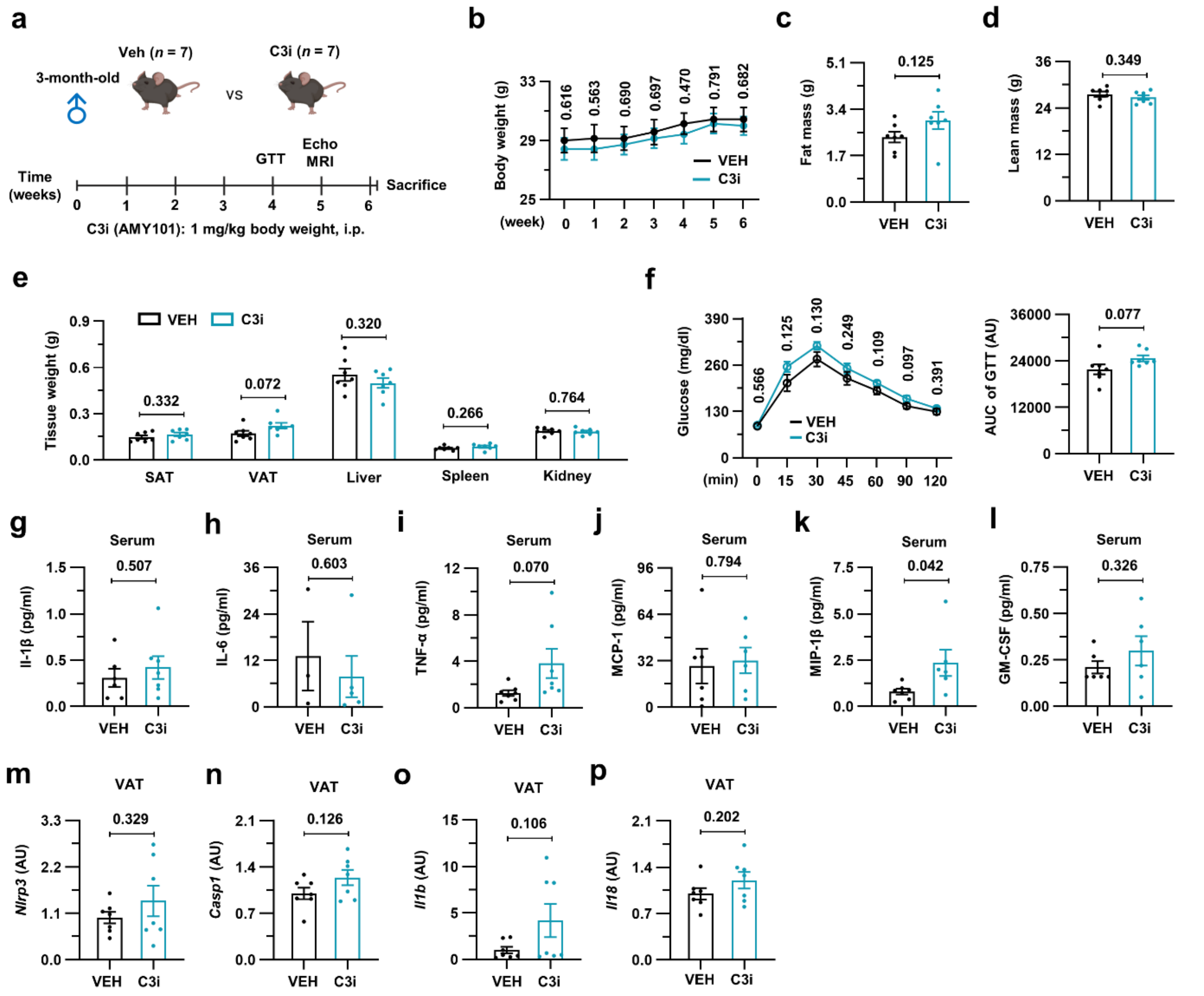


**Extended Data Fig. 6. C3 inhibitor treatment does not affect immune-metabolic homeostasis in young mice.**

(a-p) Young (3-month-old) male mice were randomly allocated and intraperitoneally injected with either vehicle or the C3 inhibitor (AMY-101; 1 mg/kg) weekly for 6 weeks and analyzed (*n* = 7/group). (a) Schematic diagram of experimental design. (b) Body weight measurement. (c,d) Fat mass (c) and lean mass (d) were measured by EchoMRI. (e) The weights of the SAT, VAT, liver, spleen, and kidney were measured. (f) A glucose tolerance test was performed. (g-l) Serum levels of IL-1β (g), IL-6 (h), TNF-α (i), MCP-1 (j), MIP-1β (k), and GM-CSF (l) were analyzed by Luminex assay. Only detected values were presented. (m-p) qRT-PCR analysis of VAT for *Nlrp3* (m), *Casp1* (n), *Il1b* (o), and *Il18* (p). Unpaired two-tailed t-tests were performed, and exact p-values were presented. Data presented as mean±SEM.

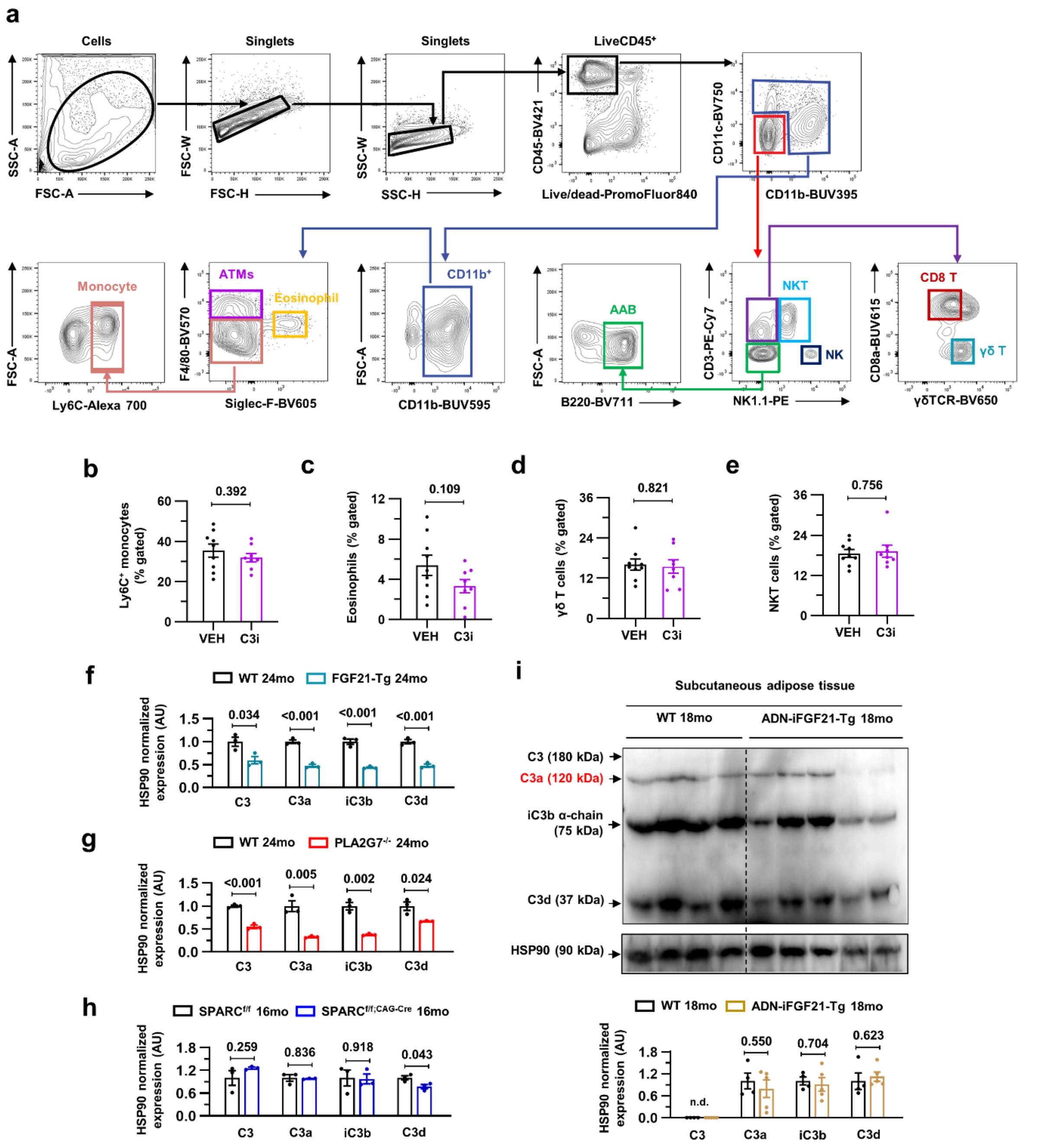


**Extended Data Fig. 7. Reduced C3 activation in mouse models of extended healthspan.** (a-e) Flow cytometry analysis of VAT immune cells in aged mice treated with either vehicle or the C3 inhibitor (AMY-101; 1 mg/kg) weekly for 6 weeks (*n* = 9 for vehicle and *n* = 8 for C3 inhibitor). (a) Gating strategy for analyzing multiple immune cell types in the VAT. (b-e) The frequencies of Ly6C$^+$ monocytes (b), eosinophils (c), γδ T cells (d), and NKT cells (e) were analyzed. (f-h) Densitometry analysis of the VAT of 24-month-old WT and FGF21-Tg (f), 24-month-old WT and PLA2G7$^{-/-}$ (g), and 16-month-old SPARC$^{f/f}$ and SPARC$^{f/f;CAG\text{-}Cre}$ (h) mice (*n* = 3/group). (i) Representative C3 western blot analysis of the SAT of 18-month-old WT (*n*

= 4) and adipocyte-specific FGF21 overexpressed (ADN-iFGF21-Tg; *n* = 5) mice with densitometry analysis. Unpaired two-tailed t-tests were performed, and exact p-values were presented. Data presented as mean±SEM.

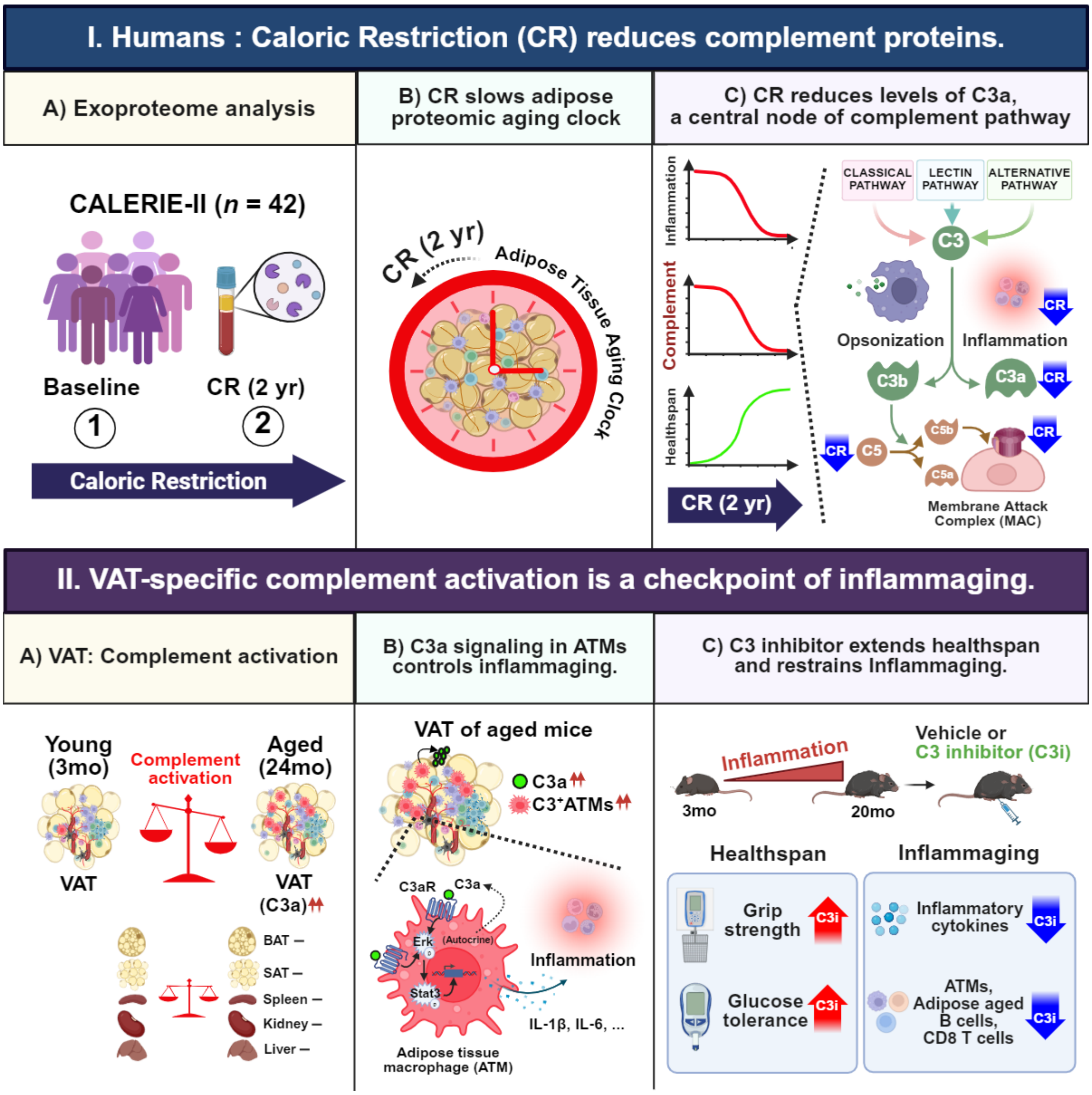


**Extended Data Fig. 8. Unbiased exoproteome analysis of calorie-restricted individuals identifies deactivation of the complement system as a checkpoint reducing inflammaging.** Longitudinal plasma proteomics analysis of CALERIE-II clinical trial participants (*n* = 42) showed that caloric restriction (CR) slows adipose proteomic aging clock, particularly suppressing the expression level of C3a, a central node of the complement pathway. The reverse translational approach revealed that visceral adipose tissue (VAT) macrophages are the major sources of age-associated increase in C3a, which promotes inflammaging in an autocrine manner in mice. The administration of clinically tested C3 inhibitor (C3i), AMY-101, demonstrated that C3 inhibition improves healthspan and restrains inflammaging in aged mice.

#### EXTENDED DATA TABLES

**Extended Data Table 1. Clinical characteristics of CALERIE trial participants for whom serum samples were used for proteomics.**

| Variables | Mean ± s.e.m. or percentage | | |
|---|---|---|---|
| | Baseline (n = 42) | CR 2 years (n = 42) | *P* value |
| Male (%) | 23.8 | | N/A |
| White (%) | 71.4 | | N/A |
| Asian (%) | 4.76 | | N/A |
| Black or African American (%) | 19.0 | | N/A |
| Age (years) | 38.29±1.22 | 40.27±1.22 | <0.001 |
| Body weight (kg) | 71.55±1.40 | 63.81±1.38 | <0.001 |
| Body mass index | 25.34±0.28 | 22.60±0.29 | <0.001 |
| Mean waist circumstance (cm) | 81.43±1.27 | 75.32±1.23 | <0.001 |
| Fat mass (kg) | 24.36±0.72 | 19.14±0.72 | <0.001 |
| Whole body fat (%) | 71.55±1.40 | 63.81±1.38 | <0.001 |
| Mean systolic blood pressure (mmHg) | 112.81±1.59 | 109.30±1.61 | 0.016 |
| Mean diastolic blood pressure (mmHg) | 73.84±1.12 | 70.4±1.21 | <0.001 |
| Serum IL-6 (pg/ml) | 1.61±0.18 | 1.43±0.29 | 0.648 |
| Serum TNF-α (pg/ml) | 3.06±0.15 | 2.57±0.14 | 0.004 |
| Serum CRP (mg/dl) | 1.59±0.43 | 0.86±0.18 | 0.048 |
| Serum leptin (ng/ml) | 20.25±2.64 | 10.44±1.46 | <0.001 |

Abbreviations: CR, caloric restriction; CRP, c-reactive protein; IL, interleukin; TNF, tumor necrosis factor.

**Extended Data Table 3. Primers for qPCR analysis.**

| Genes | Forward (5' to 3') | Reverse (5' to 3') |
|---|---|---|
| *Actb* | CTC TGG CTC CTA GCA CCA TGA AGA | GTA AAA CGC AGC TCA GTA ACA GTC GG |
| *Hprt1* | ACA GGC CAG ACT TTG TTG GA | ACT TGC GCT CAT CTT AGG CT |
| *Nlrp3* | GCT AAG AAG GAC CAG CCA GA | CAG CAA ACC CAT CCA CTC TT |
| *Casp1* | GGA CCC TCA AGT TTT GCC CT | AGA CGT GTA CGA GTG GTT GT |
| *Il1b* | GGT CAA AGG TTT GGA AGC | TGT GAA ATG CCA CCT TTT |
| *Il18* | GAC AGC CTG TGT TCG AGG AT | CAG TCT GGT CTG GGG TTC AC |
